# Dynamic alterations of DNA methylation and transcriptome in adaptation to and recovery from the space environment

**DOI:** 10.64898/2025.12.03.691098

**Authors:** Liang Lu, Yangyang Hao, Xue Lin, Kai Li, Tianzhang Zhai, Fengji Liang, Linhuan Chen, Liqing Wang, Xueyin Mei, Sidu Feng, Ke Lv, Yanhong Yuan, Zhongquan Dai, Dong Liu, Hongyu Zhang, Chao Yang, Anna Liu, Linjie Wang, Zhili Li, Shujuan Liu, Xiaoqian Dai, Chengjia Yang, Chunyan Wang, Peng Sun, Liujia Shi, Chi Zhang, Jianghui Xiong, Ming Wei, Chong Xu, Zhaoxia Liu, Lina Qu, Jian Li, Yinghui Li

## Abstract

Space exploration presents tremendous health challenges. Here, we report time series of multi-omic and phenotypic profiles of seventeen astronauts from six China Manned Space missions with continuous spaceflight durations ranging from 13 to 180 days. We revealed a key role of DNA methylation regulation in reshaping gene expression patterns to adapt to the space environment. Long-duration spaceflight showed more alterations in epigenetic modifications correlated with alternative splicing and protein acetylation. During recovery, an “overrange rebound” phenomenon was observed, furthermore, a mathematical model was established to describe this implying important phenomenon. Moreover, we revealed the correlations between molecular alterations and phenotypic changes such as coagulation activation and bone intensity loss. Additionally, we performed ground-based simulation experiments to estimate the impacts of individual stressors in the space environment on DNA methylation. In summary, our study highlights the importance and complexity of epigenetic regulation in adaptation to and recovery from the space environment.

## INTRODUCTION

With the increasing frequency of space explorations^1–3^, understanding the mechanisms of human adaptation to and recovery from space environment has become increasingly important. In space, stressors that affect human health include microgravity, radiation, confinement/isolation, and distance from Earth^4^, which collectively cause unique physiological changes in the human body^5–7^. Increasing evidence shows that most of these physiological changes are mediated by molecular events^4^. The International Space Station (ISS)^8^, which serves as a space research laboratory, has already supported many important molecular biological studies^1,9^. Over the last decade, China has launched a series of manned spacecraft Shenzhou missions^10,11^ and established the China Space Station (CSS), contributing new knowledge regarding space adaptation^12^. Here, we present analyses of multi-omic and phenotypic data from biological samples collected from seventeen astronauts who participated in the recent 10-years China Manned Space missions (M13, a 13-day spaceflight mission; M15, a 15-day spaceflight mission; M33, a 33-day spaceflight mission; M90, a 90-day spaceflight mission; M180-1, a 180-day spaceflight mission; M180-2, another 180-day spaceflight mission) (Figure 1). Bulk RNA sequencing, DNA methylation and phenotypic measurements were applied to analyze the biological samples of the astronauts mainly at three key timepoints: preflight (T1, more than 1 month before launching), immediate postflight (T2, around 1 day after returning to Earth) and recovery (T3, more than between 30 and 60 days after returning to Earth). In this study, an epigenetic characteristics of space environment adaptation and recovery on the basis of the genome-wide DNA methylation landscape from a chronological perspective was revealed. Furthermore, the results from the comparison between M90 and M180-1 missions were confirmed by the M180-2 mission, which presented the impact of long-duration spaceflight on the human body. These experiments revealed an important role of DNA methylation as a mechanism underlying human adaptation to and recovery from spaceflight on basis of data from multiple subjects and missions. Corresponding ground-based simulation experiments were also conducted^13,14^ (Figure 1 and Table S1), and these findings were compared with those from spaceflights, enabling us to investigate the impacts of individual space environmental stressors on the human body.

**Figure 1.**
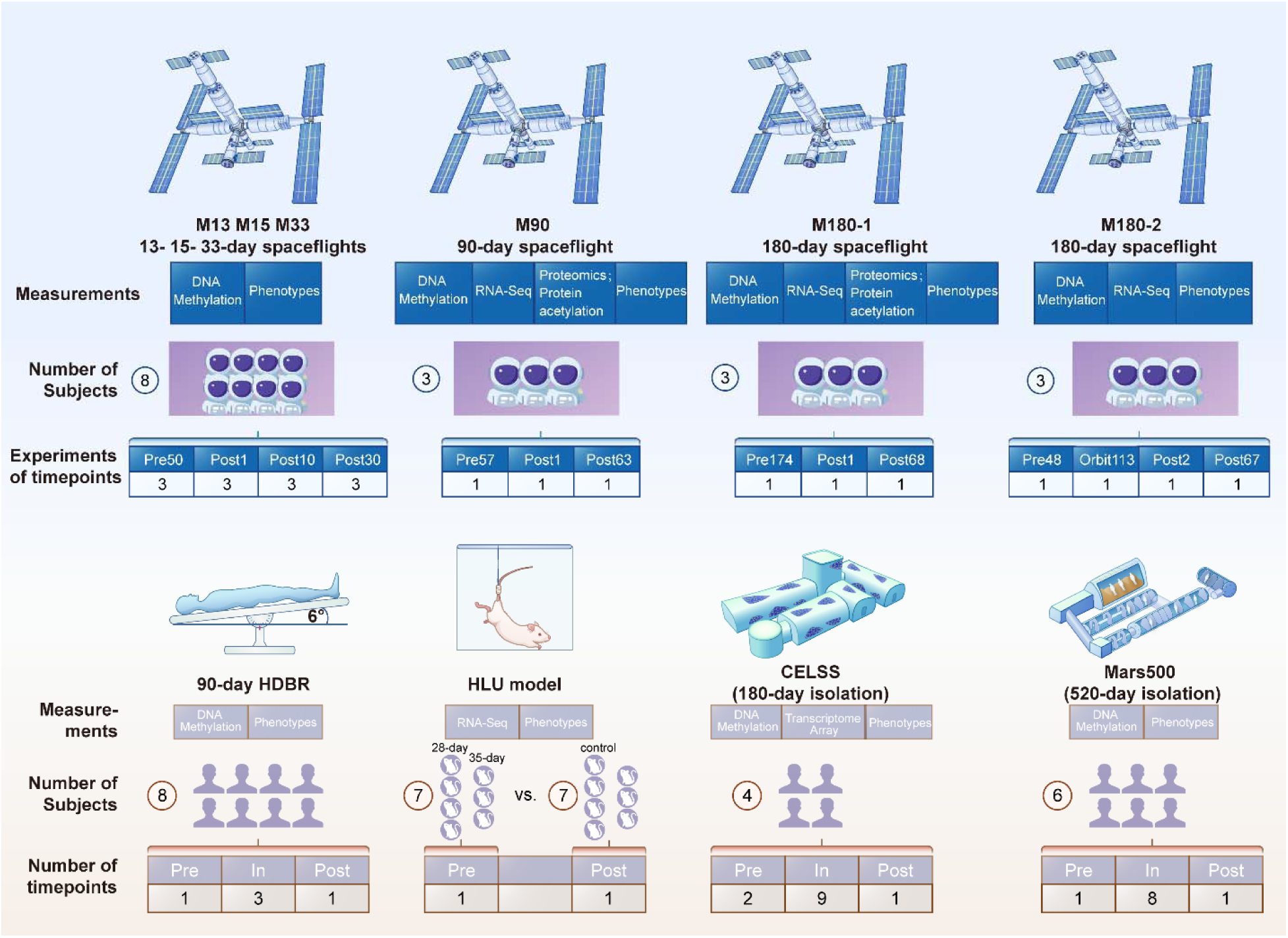
Presentation of the experimental design and data in this study We present DNA methylation and physiological (phenotypic) data from three cohorts of astronauts in the China Manned Space Program (CMSP), and transcriptomic, DNA methylation, and physiological (phenotypic) data from three cohorts of astronauts in the CSS. In addition, we performed ground-based simulations, including microgravity-stimulated head-down bed rest (HDBR) and rat hindlimb unloading (HLU), and included data from the 180-day CELSS isolation simulation and the Mars500 isolation stimulation (520 days), to elucidate space environmental stressors.

## RESULTS

### Increased DNA methylation alterations follow the accumulated space environment exposure time

We analyzed the peripheral blood DNA methylation profiles of seventeen astronauts from six China Manned Space missions, including three short-duration spaceflights (M13, M15 and M33), one medium-duration spaceflight (M90) and two long-duration spaceflights (M180-1 and M180-2). A comparison of the DNA methylation profiles of astronauts from the spaceflights with different inflight durations revealed that DNA methylation changes (T2 vs. T1) increased with increasing space environment exposure time (Figure 2a). That is, for short periods, i.e., 13 days (M13) and 15 days (M15), little change in DNA methylation was observed; a few DNA methylation alterations were identified in the 33-day spaceflight (M33); and obvious DNA methylation changes were detected after 90 days of spaceflight (M90). More significant global DNA methylation changes were present in the two 180-day spaceflights (M180-1 and M180-2). Interestingly, decreased DNA methylation was more significant than increased DNA methylation (T2 vs. T1) in almost all six spaceflights, suggesting that exposure to the space environment initiated an epigenetic transition toward more open DNA (Figure 2a). Because the six spaceflight missions included in this study span more than ten years, The DNA methylation analyses of the early spaceflight missions (M13, M15 and M33) employed the Illumina 450K platform, the analysis of the latest three spaceflights (M90, M180-1 and M180-2) employed the updated 850K platform. We used consistent probes between the 450K and 850K platforms, the general tendency that more differentially methylated probes (DMPs) were found in longer-duration spaceflights was maintained (data not shown).

**Figure 2.**
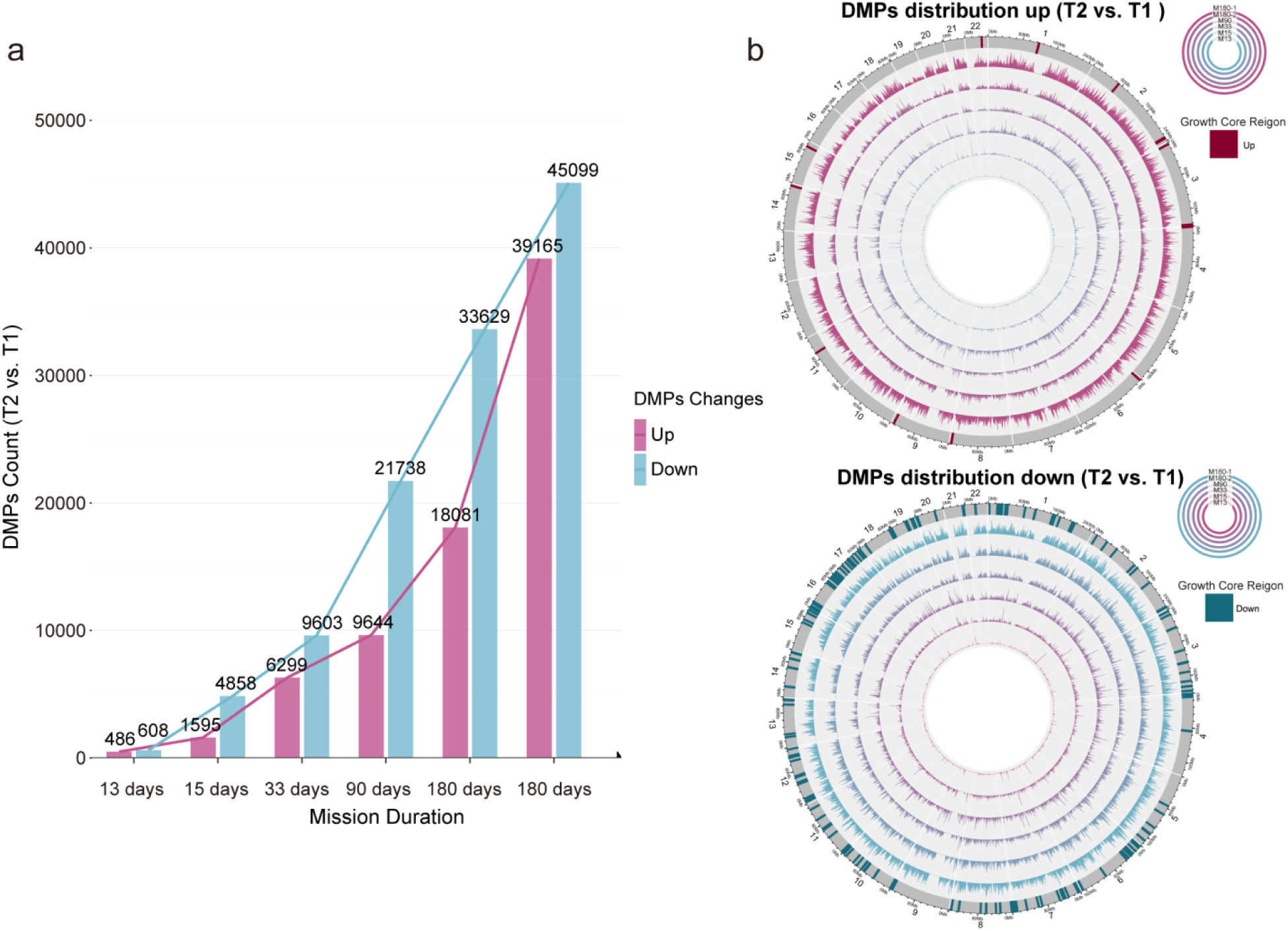
DNA methylation changes in the six spaceflights **a,** X-axis shows the six spaceflight times ranging from 13 to 180 days. The Y-axis shows the number of DMPs (T2 vs. T1), with increased methylation DMPs presented and decreased DMPs. **b,** Global changes in DNA methylation across six missions. The increased and decreased DMPs are shown above and below, respectively. The DMPs from M13, M15, M33, M90, M180-2 and M180-1 are plotted from outside to inside. Marks on the outermost circle represents the DMP enrichment region that DMPs keeps increasing following the mission duration increases, as consistent regions across all missions.

Through the comparison of the changes in DNA methylation among the six spaceflight missions, several consistent DNA methylation alteration regions were identified, which seemed to work as growth core regions. A gradual extending pattern of DNA methylation change initiated from the growth core regions was shown following increased exposure to the space environment (Figure 2b).

### Long-duration spaceflight results in increased DNA demethylation and up-regulation of gene expression

As significant changes in DNA methylation occurred during medium- and long-duration spaceflights, we further investigated the similarities and differences in DNA methylation between these two cases and their biological implications. A global change in the DNA methylation landscape occurred after spaceflight (T2 vs. T1), and most of the changes in DNA methylation were reversed during recovery, restoring the sites to their preflight status (T3 vs. T2), showing an imperfect symmetrical curve (Figure 3a and Figure S1a).

**Figure 3.**
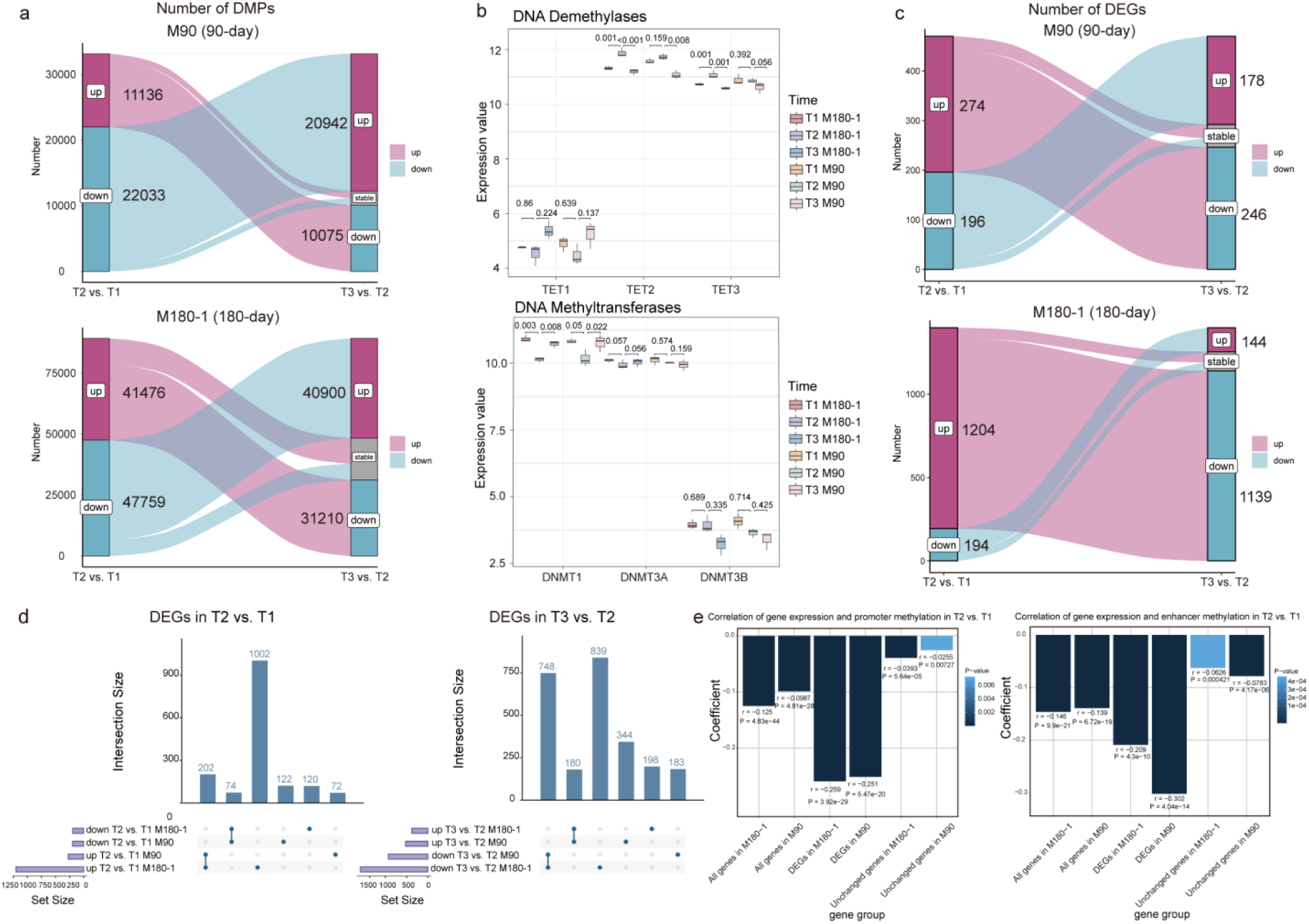
DNA methylation and transcriptome changes after spaceflight and recovery, and the correlation between DNA methylation and gene expression **a,** The top panel shows the numbers of differentially methylated positions (DMPs) after spaceflight (T2 vs. T1) and after recovery (T3 vs. T2) in the 90-day spaceflight group (M90). The bottom panel shows the same data for the 180-day spaceflight group (M180-1). **b,** Comparison of the gene expression of demethylases (top) and methyltransferases (bottom) at three timepoints (T1, T2 and T3) in M90 and M180-1. The adjusted *P* values calculated by limma are listed. **c,** The upper panel shows the number of differentially expressed genes (DEGs) after spaceflight (T2 vs. T1) and after recovery (T3 vs. T2) from the M90 group. The bottom panel shows the same data from the M180-1 group. **d,** The number of overlaps of the DEGs between the two flights (left: T2 vs. T1, right: T3 vs. T2). The dots connected below the bars represent intersections, while the separate dots represent genes unique to each group. **e,** Correlation coefficients between the log_2_FCs of the expression level and the log_2_FCs of the DNA methylation level in the promoter and enhancer regions after spaceflight (T2 vs. T1) calculated for all expressed genes, all DEGs, and unchanged genes for the two spaceflight missions.

Moreover, the number of hypomethylated DMPs was greater than the number of hypermethylated ones during spaceflight, especially for the promoter and enhancer regions (T2 vs. T1) (Supplementary Information Note S1, Figure 2a and Figure S1b), indicating genome-wide demethylation under exposure to the space environment. The above phenomenon was consistently observed in M180-2 (Figures S2a and S2b). Furthermore, longer-duration spaceflight resulted in more loci with DNA methylation changes than did medium-duration spaceflight (11136 hypermethylated / 22033 hypomethylated DMPs in M90, 41476 hypermethylated / 47759 hypomethylated DMPs in M180-1 and 16468 hypermethylated /33884 hypomethylated DMPs in M180-2, *P* value < 0.05, |deltaBeta| > 0.05, ChAMP, with consistent changes were observed among all astronauts in the same mission; see “Results of differential methylation analysis of DNA methylation probes” in https://www.spacelifescience.cn/search/detail?id=24).

The results of the differentially methylated region (DMR) analysis also support the above conclusion that M180-1 presents more DMRs of demethylation than M90 does in T2 vs. T1 (448 hypermethylated / 109 hypomethylated regions in M90, 78 hypermethylated / 976 hypomethylated regions in M180-1, adjusted *P* value < 0.05, ChAMP) (Figure S3a and Table S2). Despite a global demethylation trend indicated by DMP number in both missions and DMR number in M180-1, M90 exhibited more hypermethylated DMRs, yet only the hypomethylated DMRs were functionally cohesive enough to yield significant GO enrichment terms (Figure S3a and Table S2). Moreover, 24.1% (134) of the DMRs in M90 overlapped with the DMRs in M180-1, including 45 completely overlapping DMRs (Figure S3a and Table S2). These DMRs present in both missions were enriched for myeloid leukocyte activation, regulation of neutrophil activation and leukocyte activation involved in the inflammatory response, etc. (Table S2). The demethylated genes involved in DMRs in M90 were also predominantly immune and inflammation related genes, whereas the DMRs in M180-1 involved a greater diversity of functional gene sets, such as “locomotory behavior”, “extracellular matrix organization” and “sensory organ morphogenesis” (Table S2). Both missions present more hypermethylated DMRs than hypomethylated DMRs in T3 vs. T2 (Figure S3b). The DMRs in M180-2 still exhibited considerable overlaps with those in M180-1 and M90 (Figure S3c).

As the mammalian DNA methylome is shaped by two antagonistic processes, methylation by DNA methyltransferases (DNMTs) and demethylation by ten-eleven translocation (TET) dioxygenases^15^, we further investigated the expression of methylation-related enzymes. The upregulation of demethylases (TET2 and TET3) and the downregulation of DNMT1 were observed during spaceflight (Figure 3b and Table S3), which might be the underlying causes of the observed changes in global methylation patterns. Similar alterations were observed in M180-2 and hind-limb unloading (HLU) model rats (Supplementary Information Note S1 and Table S7).

Accordingly, longitudinal bulk RNA-seq data analysis of the astronauts’ peripheral blood samples was performed for M90 and M180-1, which revealed a dynamic global change in gene transcription with numerous differentially expressed genes (DEGs) identified (|log_2_FC| > log_2_(1.5), and adjusted *P* value < 0.05, limma). Additionally, the total number of DEGs was greater in the samples from long-duration flights than in those from medium-duration flights (Figure 3c). In the immediate postflight samples (T2 vs. T1), there were more upregulated genes (274 in M90; 1204 in M180-1) than downregulated genes (196 in M90; 194 in M180-1), revealing genome-wide significantly upregulated transcriptomic changes resulting from exposure to the space environment (Figure 3c and Table S3). This phenomenon was highly similar to that in M180-2 (Figure S4a-S4c). 202 upregulated and 74 downregulated genes overlapped in both M90 and M180-1 (T2 vs. T1), while 180 upregulated and 748 downregulated genes overlapped after recovery (T3 vs. T2) (Figure 3d).

A substantial negative correlation was observed between the changes in gene expression levels and the changes in DNA methylation levels (log_2_FCs in T2 vs. T1) in the corresponding promoter and enhancer regions for all expressed genes in both M90 and M180-1 (*P* value < 0.01, Pearson correlation test), reflecting the impact of DNA methylation on the regulation of gene expression during adaptation to the space environment (Figure 3e). This association between gene expression and DNA methylation was more prominent in the DEGs (T2 vs. T1 or T3 vs. T2) and less pronounced in the unchanged genes.

Furthermore, longer spaceflight was associated with more pronounced negative correlations (Figure 3e). The correlation between DNA methylation and gene expression observed in M90 and M180-1 was confirmed in M180-2 (Figure S4d).

### Association of differentially expressed alternative splicing isoforms with DNA methylation

Alternative splicing (AS) is a mechanism by which genes produce more than a single mature transcript, which contributes to adaptation to a new environment^16^. In our study, we found no differential AS events between the T1 and T3 transcriptomes in M90, but several differential AS events between the T1 and T2 transcriptomes (Table S4). In the comparison of AS between T2 and T1, M180-1 presented more differential AS events than M90 did (Figure 4a). However, the genes exhibiting AS in the two spaceflight durations had little overlap. The genes with differential AS events in M90 were enriched mainly in immune-related pathways, whereas the genes in M180-1 were enriched primarily in epigenetics-related pathways (Table S4), indicating that epigenetic regulation events may be enhanced during long-duration spaceflight. In M180-1, histone acetylation-related genes were found to be differentially spliced. For example, differentially expressed AS isoforms of the histone deacetylase complex subunit Sin3A-associated protein 18 (*SAP18*)^17^, as well as histone acetyltransferase-related genes *BRD1*^18^ and *JADE1*^19^, were detected after long-duration spaceflight (M180-1). In M180-2, another 180-day spaceflight, we observed differential splicing of the *MEN1* gene, which encodes the histone methyltransferase complex Menin^20^, in the comparison between T2 and T1 (Table S4), The above results suggest that multiple epigenetic regulatory mechanisms are involved in prolonged adaptation to the space environment.

**Figure 4.**
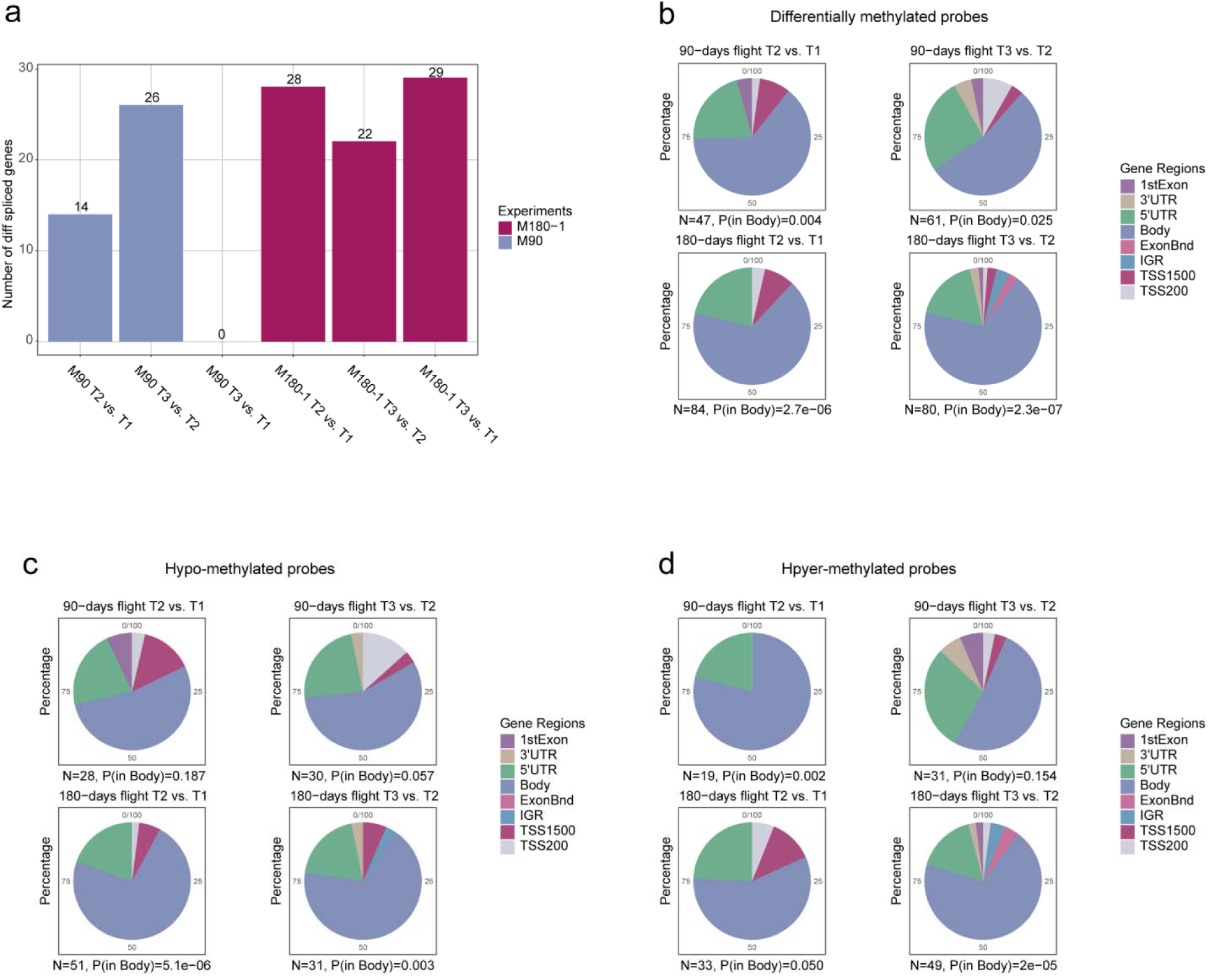
Alternative splicing events during adaptation to and recovery from spaceflight and their associations with gene body DNA methylation. **a,** The number of differential AS events between three timepoints of the two spaceflights (M90 and M180-1). **b,** The number of DMPs occurring within genes with significant AS events, categorized and counted according to the regions of the gene, on which the probes are located. A hypergeometric test was performed to evaluate the significance of enrichment of the loci in the gene body regions. **c, d,** Similar to **b** but for the number of hypomethylated and hypermethylated probes, respectively. TSS, transcriptional start site; TSS200, 0–200 bases upstream of the TSS; TSS1500, 200–1500 bases upstream of the TSS; 5’UTR, within the 5’ untranslated region, between the TSS and the ATG start site; Body, between the ATG and stop codon; irrespective of the presence of introns, exons, TSS, or promoters; 3’UTR, between the stop codon and poly A signal; ExonBnd, within 20 bases of an exon boundary (i.e., the start or end of an exon); IGR, intergenic regions.

The loci with altered DNA methylation levels in the differentially spliced genes, especially hypomethylated loci, were significantly localized in gene body regions rather than randomly distributed (Figures 4b-4d and Table S4). Previous studies have reported that DNA methylation plays a role in the identification of exons involved in AS events, and that the loss of DNA methylation affects splicing patterns on a genome-wide scale^21,22^. Accordingly, we speculate that DNA methylation may be involved in the regulation of AS events during adaptation to the space environment.

### Changes in proteomics and protein acetylation levels during spaceflight and recovery

Differential analysis of the proteomic profiles revealed 8 proteins that were differentially expressed postflight in M90 and 14 proteins that were differentially expressed postflight in M180-1 (adjusted *P* value < 0.05, limma). Reductions in hemoglobin levels were detected in astronauts from both flights (M90 and M180-1) (Table S5). It has been reported that lower hemoglobin accelerates platelet aggregation, thereby promoting coagulation^23^. The expression of Ig-like domain-containing, proteins, which are typically involved in modulating immune responses, decreased postflight in M90^24,25^. The myeloblastin protein, also known as PRTN3, showed increased expression postflight in M180-1. PRTN3 is a neutrophil-specific protease primarily associated with the neutrophil immune response and inflammatory processes^26^. The number of proteins with acetylation modifications was generally reduced after both spaceflights (T2 vs. T1) (Table S5), especially M180-1 (*P* value = 0.077, Wilcoxon test, two-tailed) (Figure S5a).

Interestingly, all proteins that were differentially acetylated exhibited ‘V’ shaped profiles, presenting decreases in acetylation during spaceflight (T2 vs. T1) and subsequent increases during recovery (T3 vs. T2) (Figure S5b and Table S5). The functions of these differentially acetylated proteins are related primarily to the cytoskeleton and metabolic functions. Among these, the deacetylation of ATP-citrate synthase may affect cholesterol metabolism^27^, which is notable as changes in blood cholesterol levels postflight have been reported^9^. A change in the acetylation level of H1e was found only in M90, which was consistent with the global acetylation profile, which presented a ‘V’ shape (Table S5). Changes in H4 histone acetylation at H4K12 and H4K16 were detected in both M90 and M180-1 but presented different patterns. In M90, acetylation decreased at T2 and T3. However, in M180-1, H4 acetylation levels at H4K12 and H4K16 increased (T2 vs. T1) and subsequently decreased (T3 vs. T2) (Table S5), suggesting a difference in H4 histone modification between medium-duration spaceflight and long-duration spaceflight.

Interestingly, transcriptomic analysis revealed that several histone deacetylase-encoding genes^28^ *HDAC4* (log_2_FC = 0.58, adjusted *P* value = 0.001, limma), *HDAC5* (log_2_FC = 0.63, adjusted *P* value = 0.003, limma) and *HDAC7* (log_2_FC = 0.51, adjusted *P* value = 0.005, limma) were significantly increased in M180-1 (T2 vs. T1) (Table S3), with a decrease in DNA methylation in the *HDAC7* promoter region. None of these transcriptomic changes were observed in M90.

### Overrange rebound of gene expression during recovery and its underlying mechanisms

Most of the genes whose expression significantly changed during spaceflight tended to recover after returning to Earth in both M90 and M180-1. A total of 89.8% of the upregulated genes and 90.8% of the downregulated genes in M90 and 94.6% of upregulated genes and 74.2% of the downregulated genes in M180-1 showed the reverse change in expression after landing, and the expression tended to return to the preflight level (T1) (Figure 3c). However, the majority of these genes did not return to precisely the same level of expression as T1 but rather exceeded the previous expression levels to a small extent, showing “overshoot” (Figure 5a). During recovery, this overshoot was observed in 72.2% of the upregulated genes and 74.7% of the downregulated genes in M90 and 73.2% of the upregulated genes and 55.5% of the downregulated genes in M180-1 (Figure 5b). We name this phenomenon “overrange rebound” and note that it presents an imperfect symmetrical curve (Figure 5c). This phenomenon was also observed in M180-2 (Figures S4a and S4c). The functions of these overrange rebound DEGs (DEG_over) vary, including roles in immunity, the inflammatory response, bone resorption, and coagulation (Table S3). Furthermore, we established a mathematical model to describe the observed overrange rebound overshoot (Supplementary Information Note S2).

**Figure 5.**
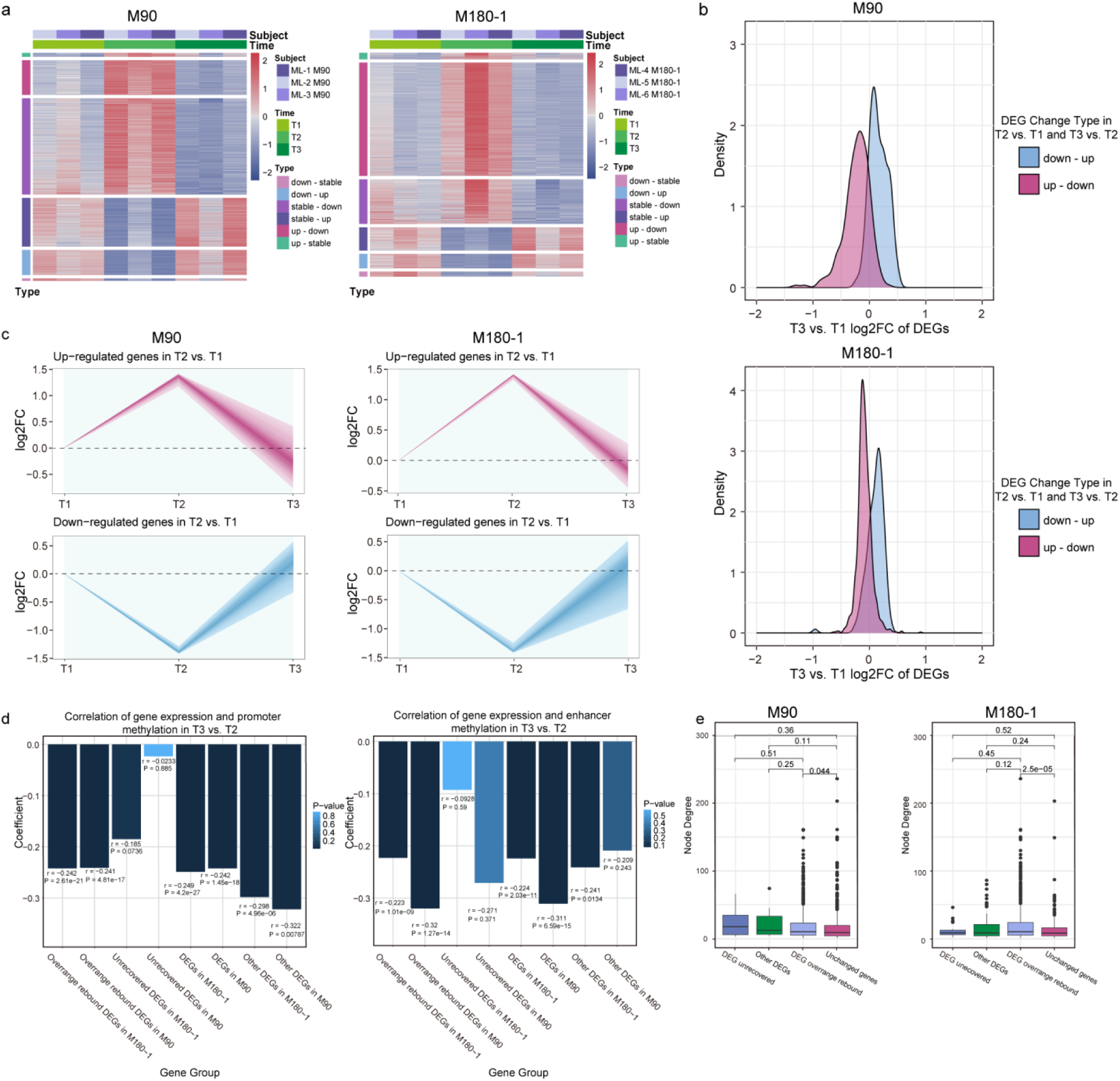
The expression of the overrange rebound genes during recovery and the effects of DNA methylation **a,** The expression profiles of all genes of all astronauts in M90 and M180-1 are shown, with genes grouped by the types of change in the T2 vs. T1 and T3 vs. T2 comparisons, e.g., “down_up” indicates downregulated in T2 vs. T1 and upregulated in T3 vs. T2, whereas “stable” indicates no significant change. **b,** Quantification of the overrange rebound of gene expression during recovery by calculating the log_2_FCs comparing T3 and T1 for the DEGs in M90 (top) and M180-1 (bottom). **c,** Dynamic changes in the expression levels of DEGs from T2 vs. T1 postflight (T2) and recovery (T3) in M90 (left) and M180-1 (right). **d,** Correlation coefficients between the log_2_FCs of the expression level and the log_2_FCs of the DNA methylation level in the promoter and enhancer regions (T3 vs. T2) calculated for all DEGs, unrecovered DEGs, over-range rebound DEGs and other DEGs in M90 and M180-1. **e,** The node degrees of various gene groups within the PPI network (Wilcoxon test, two-tailed), including DEGs (categorized as unrecovered DEGs, overrange rebound DEGs and other DEGs) and unchanged genes, in M90 (left) and M180-1 (right).

Furthermore, we found that the expression values of the overrange rebound DEGs were significantly negatively correlated with their promoter and enhancer regions DNA methylation levels (*P* value < 0.01, Pearson correlation test), moreover this correlation was not significant for the unrecovered DEGs (Figure 5d). This finding suggests a role for DNA methylation in recovery. In addition, the overrange rebound genes exhibited greater degree values in the constructed protein-protein interaction (PPI) network (“Construction of PPI networks related to environmental adaptation” in Methods) than non-rebound overrange genes did (*P* value = 0.044 for M90 and *P* value = 2.5e-05 for M180-1, Wilcoxon test, two-tailed) (Figure 5e), indicating that they interact with more genes and may be more susceptible to cascade and feedback mechanisms. Overrange rebound overshoot may be a mechanism, by which the human body can overcome the inertia established during adaptation to the space environment and accomplish rapid recovery upon return to Earth.

### Interactions of adaptation-related genes suggest that inflammation-like mechanisms coordinate multisystem adaptation

Weighted gene coexpression network analysis (WGCNA) revealed genes whose expression exhibited a characteristic chronological pattern according to the course of adaptation and recovery during spaceflight (M90 and M180-1). These genes were categorized into 7 modules on the basis of their expression correlation (Table S6), with the blue and brown modules significantly associated with spaceflight time (*P* value < 0.05, ANOVA test) (Figure 6a and Table S6). Functional enrichment analysis revealed that the genes in blue module were related mainly to immune system function, whereas the genes in brown module were involved in multiple systems of interest (Figures 6b and 6c and Table S6), including immune, bone resorption, coagulation, and oxidative stress-related gene sets. A predicted space-adaptation-related PPI network was constructed on the basis of the genes in the brown module, which reflects potential multisystemic changes induced by the space environment (Table S6). A gene set interaction network was produced by calculating the interaction score for each pair of gene sets in the 476 biological processes enriched among the 1458 PPI nodes (Table S6). Inflammation-related pathways, such as the chemokine, TNF signaling and Toll-like signaling pathways, interacted the most strongly with multisystem-related pathways (Figure 6d and Table S6). Multisystem phenotypic changes induced by the space environmental exposure, such as bone loss and blood coagulation activation, were identified in M90 and M180-1 (Table S6).

**Figure 6.**
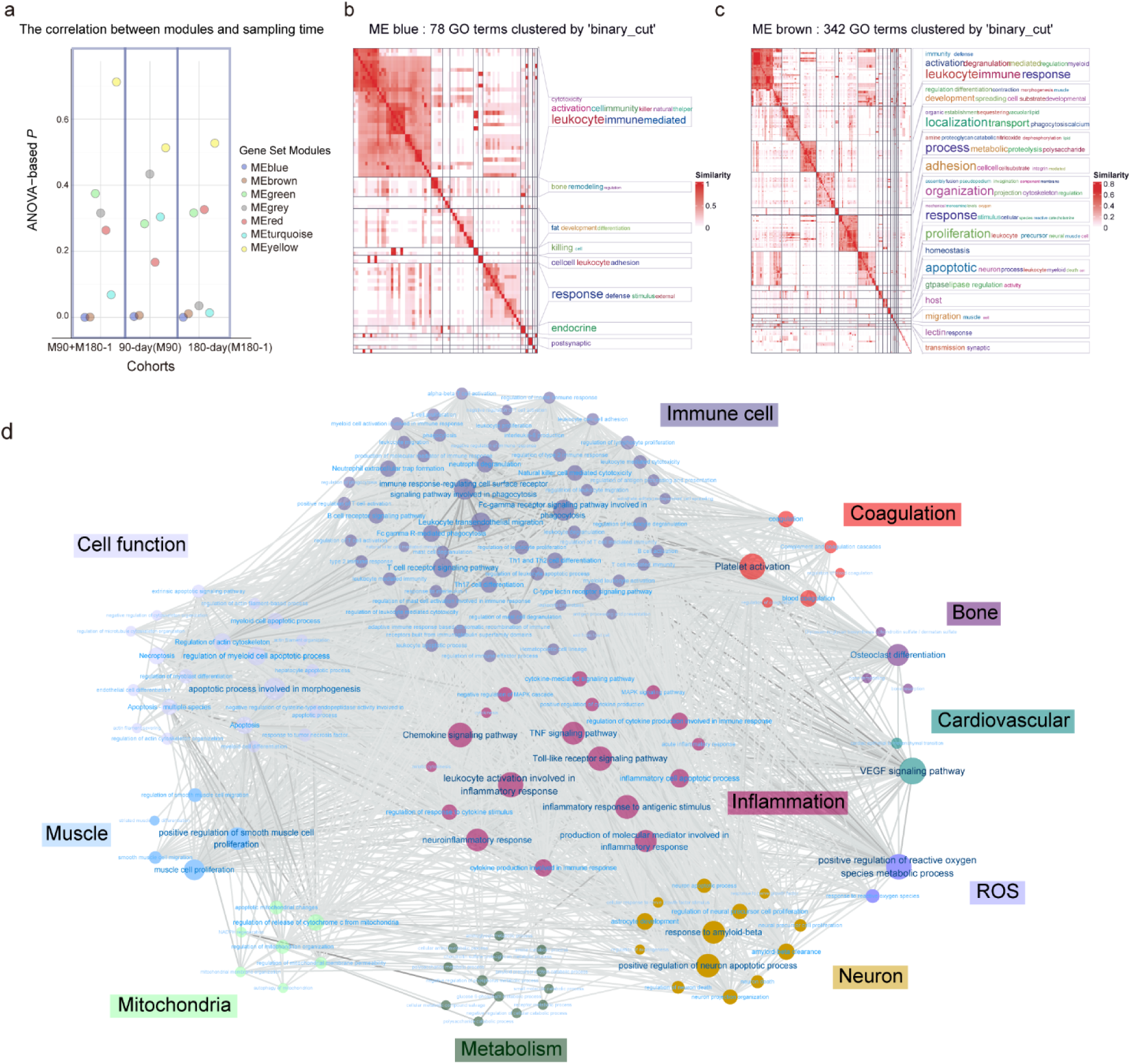
Functional enrichment results for the genes included in various modules and the PPI network constructed from the brown module. **a,** Significance of the correlation between the gene modules identified by WGCNA and time. Correlations were calculated separately in both spaceflights and jointly by ANOVA between module scores and sampling times. **b, c,** GO terms enriched among blue (**b**) and brown module (**c**) genes after clustering; keywords in different categories are displayed in descending order of occurrence. **d,** Interactions between gene sets enriched among the PPI network genes. Edges represent significant interactions, with darker to lighter colors corresponding to the intensity of interactions, and dots represent gene sets, which have been classified by function and marked with different colors.

The bone density of the six astronauts (M90 and M180-1) decreased to some extent after the spaceflights, according to phenotypic measurements, including ultrasonic calcaneal bone density, lumbar spine bone density, and left and right hip bone density. The 28-day HLU model experiment also revealed a reduction in bone density (Supplementary Information Note S3, Figure S6, and Table S7). For astronauts, the functions “osteoclast differentiation” and “NF-κB signaling pathway” were enriched in the PPI network of the brown module and generally exhibited upregulated gene expression (Table S6).

Osteoclasts are specific cells responsible for bone matrix degradation^29^. Moreover, the expression of these two gene sets was significantly negatively correlated with bone density (Table S6). Furthermore, these genes displayed a general upregulation of expression, with some experiencing decreased DNA methylation in the promoter region and overrange rebound.

The interaction of nuclear factor-kappa B receptor activator (RANK) with its ligand RANKL activates a major signaling pathway for osteoclast formation, which regulates adequate osteoclast differentiation through activation of NF-κB and its downstream targets, and genes related to these molecules were generally upregulated after both spaceflights (Table S6). In addition, the formation of mature osteoclasts may require costimulatory signals involving immune receptors associated with ITAM-carrying aptamers such as FcRγ and DAP12, most of which were significantly upregulated after the two spaceflights (Table S6). We also observed the upregulation of certain inflammatory factors, such as TLR-4^30,31^ and IL1B^32^ (Table S3), that can stimulate the NF-kappa B signaling pathway^33^. The proinflammatory cytokine *IL1B* was significantly coexpressed with the NF-κB encoding genes *NFKB1* (*r* = 0.533, *P* value = 0.020), *NFKB2* (*r* = 0.796, *P* value = 7.829e-05), and *RELB* (*r* = 0.873, *P* value = 2.241e-06) in the 18 samples from the spaceflight durations. Moreover, the range of upregulation of the above genes was generally greater in M180-1 than in M90 (Table S3). Additionally, a greater number of genes in M180-1 exhibited decreased DNA methylation and overrange rebound (Table S6).

The functions “blood coagulation” and “platelet activation” were also enriched in the PPI network of the brown module (Table S6). Two of the astronauts presented increased levels of thrombin antithrombin complex (TAT) in M180-1, indicating activation of the coagulation system^34^. Moreover, all three astronauts presented an increase in CD62P (P-selectin) after 180 days of spaceflight, indicating that platelet activation may contribute to coagulation activation^35,36^. Moreover, the expression of “blood coagulation” and “platelet activation” genes was positively correlated with CD62P and TAT. Similar phenotypic changes were detected in the 28-day HLU model rats (Supplementary Information Note S3). Gene set interaction analysis revealed that blood coagulation interacted significantly with platelet activation (Table S6). Platelets carry a wide range of chemokines and cytokines and play important roles in a variety of processes, such as hemostasis, wound repair, and pro- and anti-inflammatory activities^37^. This study revealed a significant interaction between platelet activation and chemokines and cytokines signaling associated pathways (Table S6). The expression levels of the chemokines *CXCL1* and *PF4*^38^ increased after both spaceflights (Table S3). *PF4* expression was also increased in 28-day HLU model rats.

Hemostasis-related genes encoding a vascular hemophilia factor, von Willegrand factor (vWF) were also up-regulated after 180 days of spaceflight (Table S3). vWF interacts with collagen-GPVI, initiating a series of signal transduction molecules and leading to platelet activation^39^. *TBXAS1*, αIIbβ3 integrin, and *ITGA2B*, which encodes the coagulation-related protein TXA2 synthase^40^, were also found to be up-regulated (Table S3), which could serve as evidence of coagulation activation^41,42^. Inflammation and coagulation are two major host defense systems that mutually interact^43,44^. The inflammatory cytokine IL-1β may facilitate increased expression of COX-2^45,46^ (*IL-1B* and *PTGS2*, *r* = 0.778, *P* value = 0.0001), which in turn is linked to increased expression of *TBXAS1*. In addition, we detected elevated postflight expression of the coagulation factor V encoding gene *F5* after both spaceflights (Table S3); coagulation factor V is an important cofactor in the blood coagulation cascade. Overall, there was greater up-regulation of coagulation-related genes in M180-1 than in M90, which may explain the coagulation activation observed after 180 days of spaceflight. In addition, more genes in M180-1 showed decreased DNA methylation and overrange rebound (Table S6).

In M180-2, a general postflight decrease in bone density and an increase in TAT were observed, which is consistent with the findings from the M90 and M180-1 missions. Additionally, the plasma prothrombin time (PT) and activated partial thromboplastin time (APTT) were reduced, further supporting the activation of coagulation. However, the level of CD62P decreased. The above important functional gene set also showed an increasing trend in expression in M180-2 and in HLU model rats, although individual variations were observed.

We hypothesized that, during adaptation to and recovery from the space environment, many inflammation-related genes and pathways are recruited to coordinate different organs and systems to disrupt previous physiological homeostasis and rebuild novel fitness homeostasis as a fast response to environmental change.

### Comparison of ground-based simulation experiments and real spaceflight

Because of the key role of DNA methylation in human adaptation to and recovery from the space environment revealed by the above findings from real spaceflights, we further investigated the DNA methylation patterns of different components of space environmental stressors via ground-based experiments. We compared the DNA methylation profiles of samples from M90 and M180-1 with those of samples from 90-day head-down bed rest (HDBR) (simulating microgravity), 180-day Controlled Ecological Life Support System (CELSS) (simulating 180-day isolation)^13^, and Mars500 (simulating 520-day isolation)^14^ experiments. Overall, global DNA methylation differences were detected in the 90-day HDBR and the Mars500 groups, whereas no significant global DNA methylation changes were detected in the 180-day CELSS (Figures S7a-S7c) group. The group that experienced 90 days of HDBR (R90) clearly presented significant differences compared with the prebed rest group (Pre15). A portion of samples at day 60 of HDBR (R60) presented a change in DNA methylation levels, whereas samples from day 30 exhibited little effect (Figure S7a). The DNA methylation data from Mars500 indicated notable differences in samples taken before the 60th day compared with samples taken after the 120th day, and especially samples from the 512th day (Figure S7c).

In all the aforementioned simulation studies, changes in bone phenotypes were observed. Among the eight 90-day HDBR subjects, more than six presented decreased bone densities in the hip, tibia, and calcaneus at R60 and R90 (Figure S7d). In the 180-day CELSS study, all four subjects showed a trend toward reduced bone density in both weight-bearing bones (lumbar spine and femur) and non-weight-bearing bones (radius and ulna)^47^. The Mars500 study indicated that prolonged confinement in closed environments can lead to localized reductions in bone mineralization^48^. However, these scenarios, unlike actual spaceflight and HLU models, induced localized decreases in bone density rather than a significant overall reduction in mineral density. No direct evidence of coagulation activation was found in the simulation studies. In the 90-day HDBR samples, the ADP-induced platelet aggregation rates decreased and the PAC1 levels increased at R30 and R60, but the levels of CD62P and other markers did not change significantly (Figure S7e). A previous 60-day HDBR study reported that 60 days of 6° HDBR did not correlate with significant activation of the coagulation system as evaluated via intravascular thrombus formation in healthy volunteers^49^. In the 180-day CELSS study, an increase in PAC1 was observed, but CD62P did not exhibit consistent changes (Figure S7f). The 520-day isolation study (Mars500) reported a slowdown in the intrinsic pathway and final-stage blood coagulation, with a relative increase in the activity of anticoagulant proteins such as antithrombin III (ATIII) and protein C (PC) inhibitors^50^.

Furthermore, our findings revealed that most genes related to osteoclast differentiation and the NF-κB signaling pathway presented increased expression and decreased DNA methylation (T2 vs. T1) during both M90 and M180-1 (Table S6 and Figures 7a and 7b). Among the ground-based experiments, the DNA methylation decreased in the 90-day HDBR experiment but the decrease was minimal and reverted quickly (Figure 7c). DNA methylation also decreased in the 180-day CELSS experiment on the 90th and 150th days but was not maintained (Figure 7d), and many of these genes were temporarily upregulated in the 30- and 90-day samples (Table S6). Multiple fluctuating in DNA methylation also occurred during the Mars500 mission (the 168th and 300th days) (Figure 7e). For the NF-κB signaling pathway, the fluctuations in DNA methylation did not decrease after 90 days in either HDBR or 180-day CELSS, suggesting that the effects of these experiments were not sufficient to induce stable modifications at the DNA methylation level (Figures 7c-7e).

**Figure 7.**
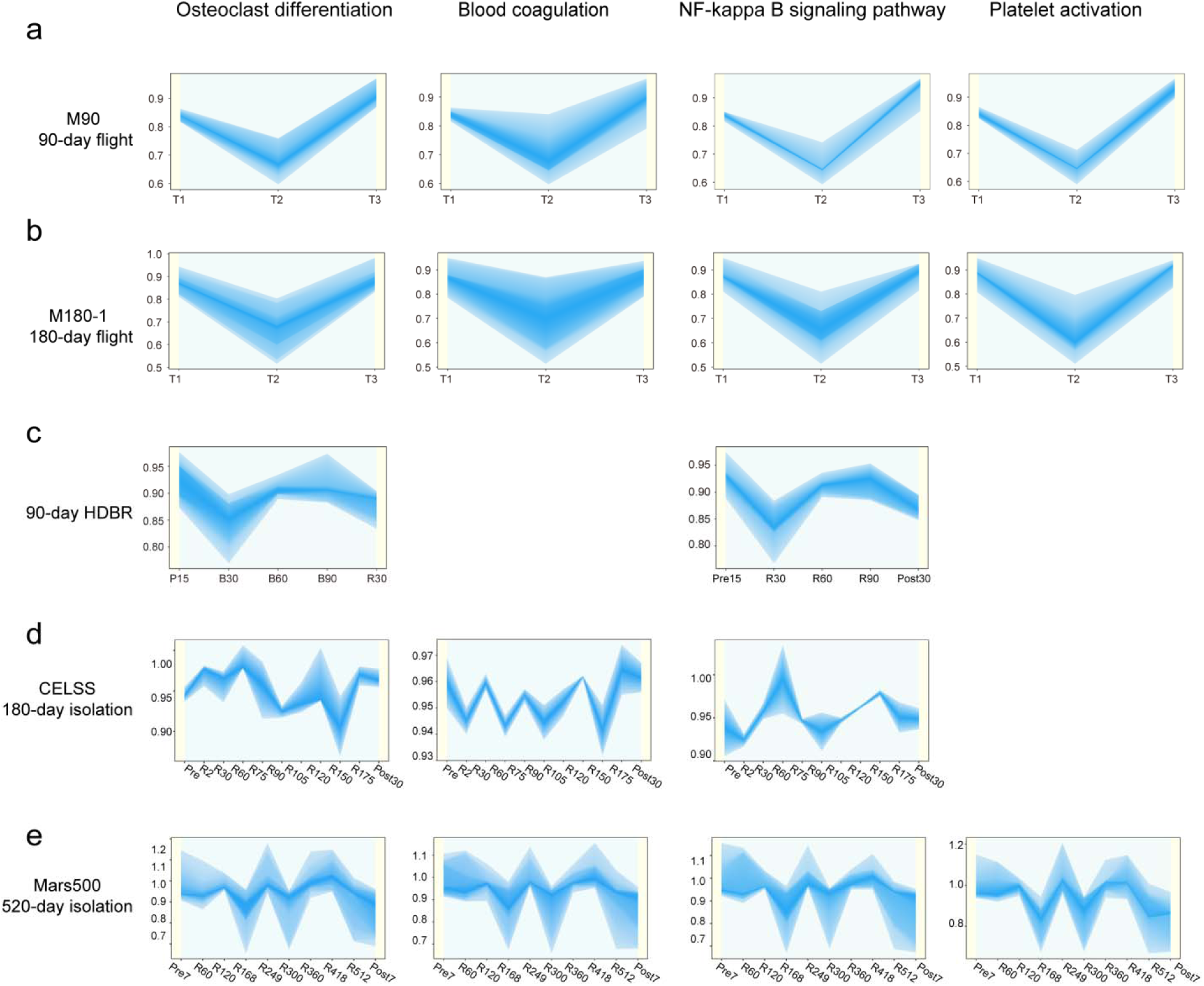
Comparison of DNA methylation changes in the gene sets associated with space environment adaptation in spaceflight and ground-based experiments. **a-e,** DNA methylation alterations of the PPI network genes enriched in the “osteoclast differentiation”, “NF-κB signaling pathway”, “blood coagulation”, and “platelet activation” gene sets in M90 (**a**), M180-1 (**b**), the 90-day HDBR (**c**), the 180-day CELSS (**d**), and Mars500 (**e**).

The DNA methylation levels of the coagulation gene set and platelet activation were not consistently altered in the 90-day HDBR and 180-day CELSS studies, and only transient fluctuations were observed in the Mars500 mission (Figures 7c-7e).

The range of DNA methylation fluctuations in the ground-based simulated experiments was generally narrower than those in actual spaceflight, with the largest fluctuations occurring during the Mars500 mission. However, the effect of time on the range of fluctuations in the Mars500 mission experiment was not found.

## DISCUSSION

The NASA Twins Study was the first to systemically reveal the molecular profiles of long-duration spaceflights on the basis of multi-omics data collected from an astronaut and his ground-based identical twin^9^. In this study, we revealed a clear dynamic molecular process occurring during the course of adaptation to and recovery from the space environment on the basis of molecular and phenotypic data from seventeen astronauts from the six China Manned Space missions. Crucially, widespread promoter DNA demethylation correlates with upregulation of gene expression, especially during long-duration spaceflight, which may cause significant upregulation of many genes, suggesting a nonnegligible role for DNA methylation in the adaptation of the human body to the space environment. With accumulated space environment exposure time, DNA methylation alterations initiated at certain core seed loci gradually extended to neighboring genomic regions to expand an open epigenetic state, reshaping the transcriptome to adapt to the space environment. With increasing spaceflight time, in addition to the gradual expansion of DNA methylation alterations from a few loci to the full genome, suggesting that multiple steps and fine turning epigenetic regulations are involved in adaptation to the space environment.

Interestingly, numerous genes were found to use different transcript isoforms during spaceflight, which indicates that AS may contribute to adaptation to the space environment. Notably, the genes with AS in M180-1 and M180-2 are involved in histone acetylation. These findings indicate that long-duration spaceflight results in more changes in epigenetic modifications than medium-duration spaceflight does and suggest that cross-talk between multiple epigenetic modifications. These epigenetic modifications such as DNA methylation and protein acetylation, orchestrate transcriptome reshaping adaptation to long-term space environmental exposure. Our study revealed that epigenetic regulatory mechanisms may determine the plasticity and elasticity of the human body in the context of rapid and long-term environmental changes.

An overrange rebound during recovery was observed in multiple cohorts, which may drive the rapid recovery upon return to Earth. Overrange rebound allows the system to respond more quickly to changes or perturbations and can help the system overcome any resistance or inertia that might be present in the system to achieve a more stable and well-regulated state in the long term. This phenomenon can be a feature of adaptive systems, in which the system learns from its past experiences and adjusts its response accordingly to refine its responses and adapt to changing environmental conditions. Our data provide some insights into the regulatory mechanism of overrange rebound by revealing the relationship between DNA methylation changes and gene expression. Further investigation is necessary to more fully comprehend the mechanisms involved, which may be more complex.

Gene expression is correlated with phenotype, and a time effect dependent correlation was found between gene sets associated with immunity and inflammation and those associated with space environment-induced phenotypes such as bone loss and coagulation activation. Our findings highlight that inflammation-like mechanisms at the physiological level connect role multiple systems in the spaceflight adaptation and recovery. Silveira et al reported that mitochondrial stress, as a central biological hub, drives the health risks of spaceflight^6^. As mitochondria are central hubs of the immune and inflammatory system^51^, our findings are consistent with those of Silveira’s study.

Notably, renewed interest in inflammation biology has been driven by the realization that inflammation is involved in a surprisingly broad range of biological processes, including tissue remodeling, metabolism, thermogenesis, and nervous system function^52^. In the context of spaceflight, the human body recruits molecular mediators that are enriched in canonical inflammation pathways, which are connected with the stress response, metabolism and other biological processes and coordinate with major organs in the body to quickly establish a physiological adaptation state and eliminate the source of perturbations from the space environment.

Ground-based simulation experiments provide the opportunity to simulate different components of space environmental stressors^53–55^. A series of systemic comparisons with samples from multiple spaceflights revealed that the range of DNA methylation fluctuations in the simulation experiments was narrower than that in actual spaceflight, suggesting that ground-based experiments may need to be carried out longer to replicate the effects observed in space. The Mars500 mission and 180-day CELSS experiments revealed little effect of time on the degree of methylation changes, which might be attributed to the ability of the human body to recover to some extent during the isolation experiment. Conversely, the effect of time in spaceflight suggests that DNA methylation changes in these gene sets may not be recoverable within the space environment. Understanding the molecular mechanisms of dynamic space adaptation necessitates the integration of multi-omics signals from multiple individuals and missions. Nevertheless, our study reveals that DNA methylation plays an important role in space adaptation and recovery, providing new insights in space life science and directions for measures to mitigate the health risks associated with spaceflight.

### Limitations of the study

Considering the limitations of actual spaceflight conditions, our study inevitably has several limitations. First, proteomics and acetylation modification analyses were performed only for M90 and M180-1 because of the limited number of materials available. Further proteomics analyses are planned for future spaceflights to investigate the roles of proteins and their modifications in human space adaptation. Second, DNA methylation and the transcriptome were analyzed via bulk sequencing; thus, the potential influence of changes in cell composition on these observations could not be eliminated. In the future, cell sorting or single-cell sequencing could be applied to evaluate the cell composition at the single-cell level. Finally, as a common limitation of spaceflight studies, few subjects are available. However, as spaceflights and space exploration are becoming more frequent, and more accumulated data and inspiring insights will allow space life science to thrive.

## RESOURCE AVAILABILITY

### Lead contact

- Requests for further information and resources should be directed to and will be fulfilled by the lead contact, Yinghui Li.

### Materials availability

This study did not generate new unique reagents. All the reagents in this study were included in the key resources table.

### Data and code availability

- All omics data have been deposited at GSA, GSA-Human and OMIX (The Genome Sequence Archive (GSA), https://ngdc.cncb.ac.cn/gsa/; GSA-Human, https://ngdc.cncb.ac.cn/gsa-human/; OMIX, https://ngdc.cncb.ac.cn/omix/). Identifiers of the datasets are documented in https://www.spacelifescience.cn/datasets. They are available upon request if access is granted. To request access, contact the corresponding author. In addition, processed datasets derived from these data have been deposited at LSOS (https://www.spacelifescience.cn) and are publicly available as of the date of publication.
- All original code has been deposited at https://github.com/lijian2014seu/LSOS and is publicly available as of the date of publication.
- Any additional information required to reanalyze the data reported in this paper is available from the lead contact upon request.

## ACKNOWLEDGMENTS

This work was supported by the CSS Program (HYZHXMH01001, HYZHXM01004), the National Key R&D Program of China (2022YFA1104302, 2022YFA1604503), grants from the National Key Laboratory of Space Medicine, China Astronaut Research and Training Center (SMFA17B07, SMFA19A03, SMFA19B01, SMFA19C01, SMFA20C03), the National Natural Science Foundation of China (32270607, 31900473, 31800707), the Young Elite Scientists Sponsorship Program by CAST (2018QNRC001), and State Key Laboratory of Space Medicine, China Astronaut Research and Training Center (SKL2024K03, SKL2024Y04, SKL2024Y07). The authors would like to thank the astronauts for their support of this work.

## AUTHOR CONTRIBUTIONS

Conceptualization, project administration and supervision: Y.H.L., J.L., and L.N.Q.; Investigation: L.L., Y.Y.H, X.L., K.L., T.Z.Z., F.J.L., L.H.C., L.Q.W., X.Y.M., S.D.F., K.L., Y.H.Y., Z.Q.D., D.L., H.Y.Z., C.Y., A.A.L., L.J.W., Z.L.L., S.J.L., X.Q.D., C.J.Y., C.Y.W., P.S., L.J.S., C.Z., J.H.X., M.W., C.X., Z.X.L., Q.L.N., J.L., and H.Y.L.; Writing – original draft: Y.Y.H.; Writing – review and editing: J.L., X.L., L.L., Y.H.L., and L.N.Q. Funding acquisition: Y.H.L., J.L., L.N.Q., L.L., K.L., Y.H.Y, and X.L.

## DECLARATION OF GENERATIVE AI AND AI-ASSISTED TECHNOLOGIES

The authors did not use any generative AI and AI-assisted technologies in this study.

## SUPPLEMENTAL INFORMATION

**Document S1. Tables S1, FigureS1-S7,** supplemental methods, supplemental notes, and supplemental references.

**Table S2.** Results of quantitative and functional enrichment analysis of DMRs.

**Table S3.** Gene differential expression analysis results.

**Table S4.** Differentially spliced genes.

**Table S5.** Differential expression analysis results of peptides and protein acetylation modification changes in mission M90 and M180-1.

**Table S6.** PPI network.

**Table S7.** HLU model rat.

---

**Figure S1.**
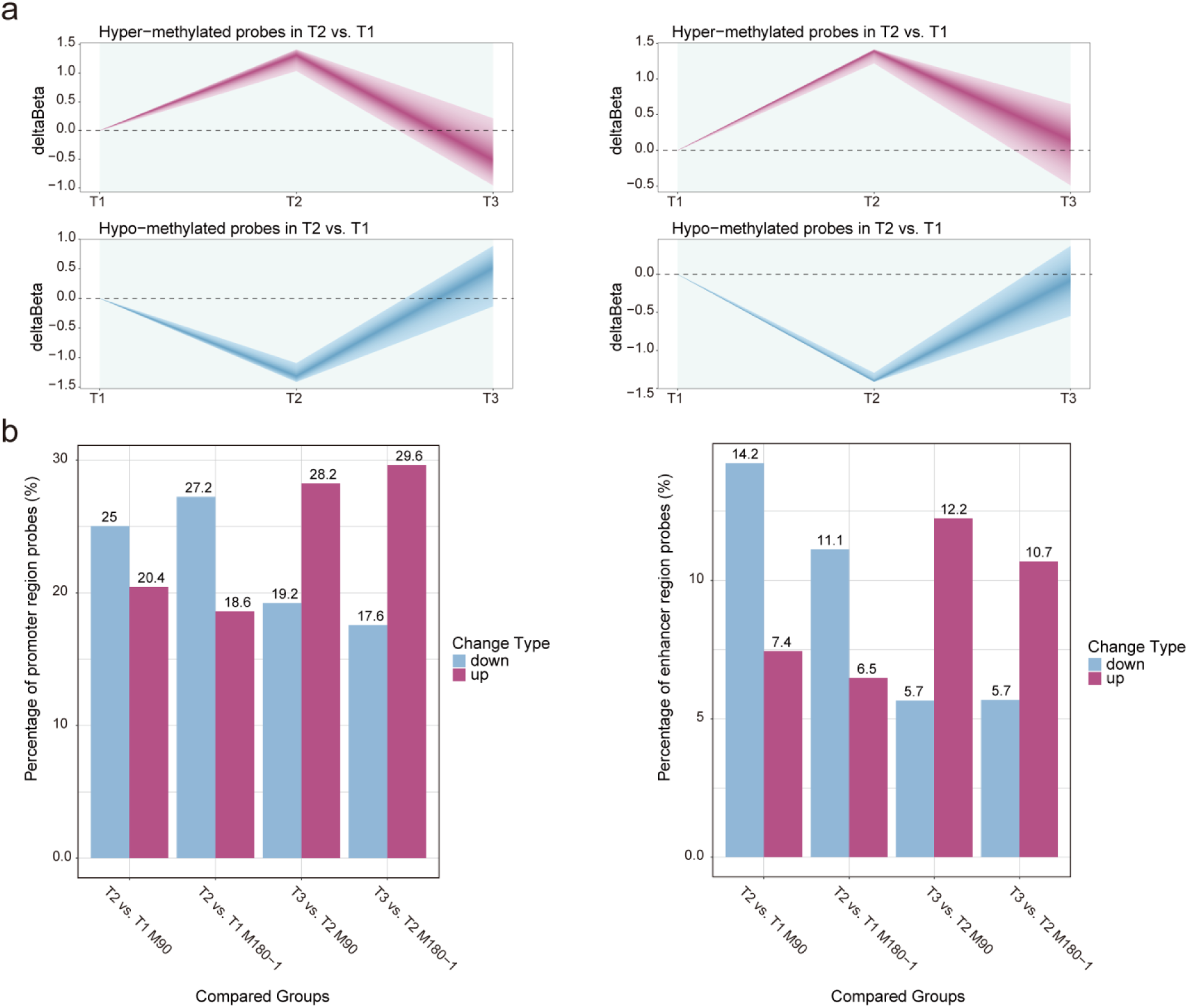
Global landscape of DNA methylation alterations in M90 and M180-1 **a,** Dynamic changes in the methylation levels of DMPs in T2 vs. T1 postflight and recovery in M90 (left) and M180-1 (right). **b,** Proportion of demethylated, hypermethylated DMPs distributed in promoter and enhancer regions compared with total demethylated, hypermethylated DMPs in each experiment.

**Figure S2.**
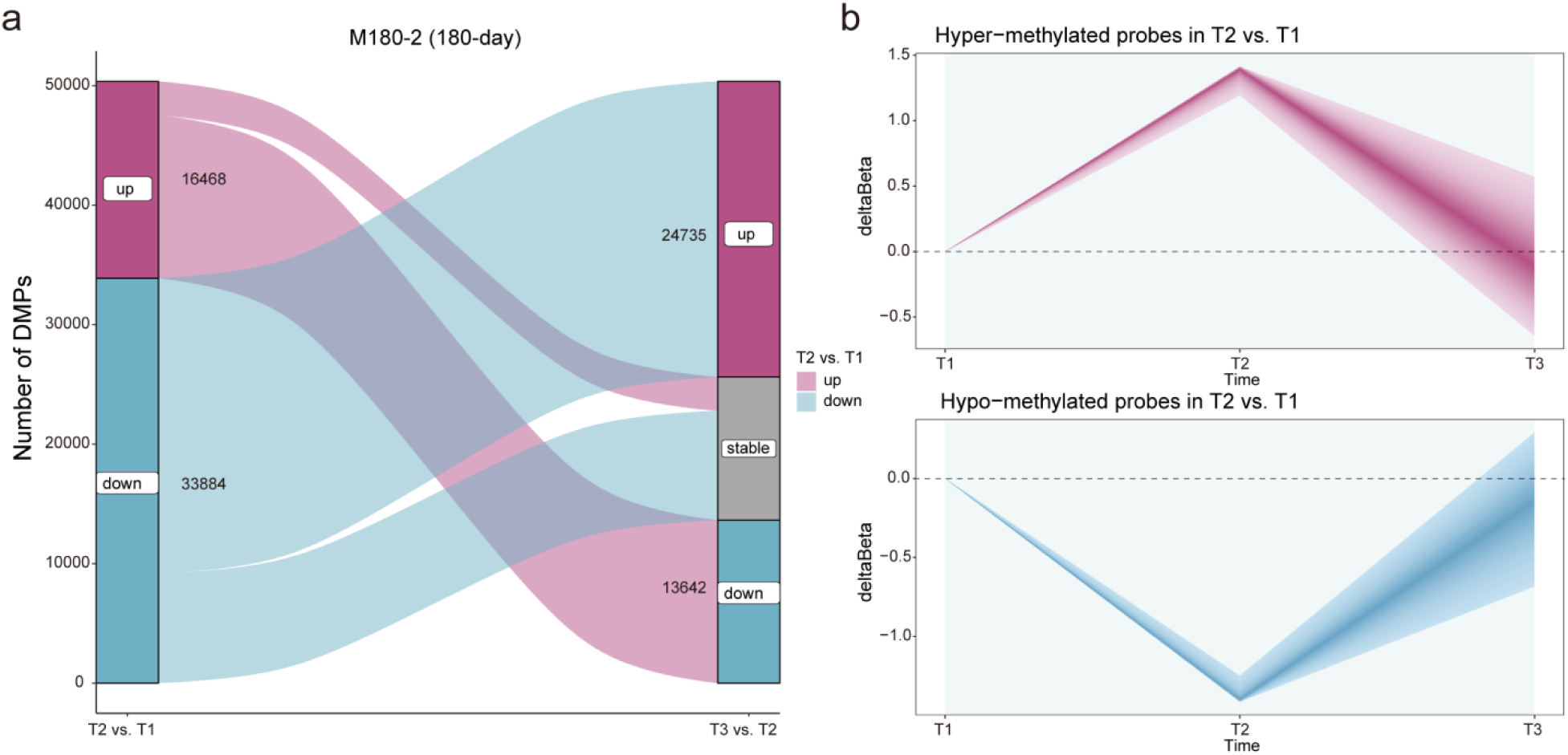
Alterations in DNA methylation during M180-2 spaceflight **a,** Number of DMPs postflight (T2 vs. T1) and during recovery (T3 vs. T2) in M180-2. **b,** Dynamic changes in the methylation levels of DMPs (T2 vs. T1) postflight and during recovery (T3 vs. T2) in M180-2.

**Figure S3.**
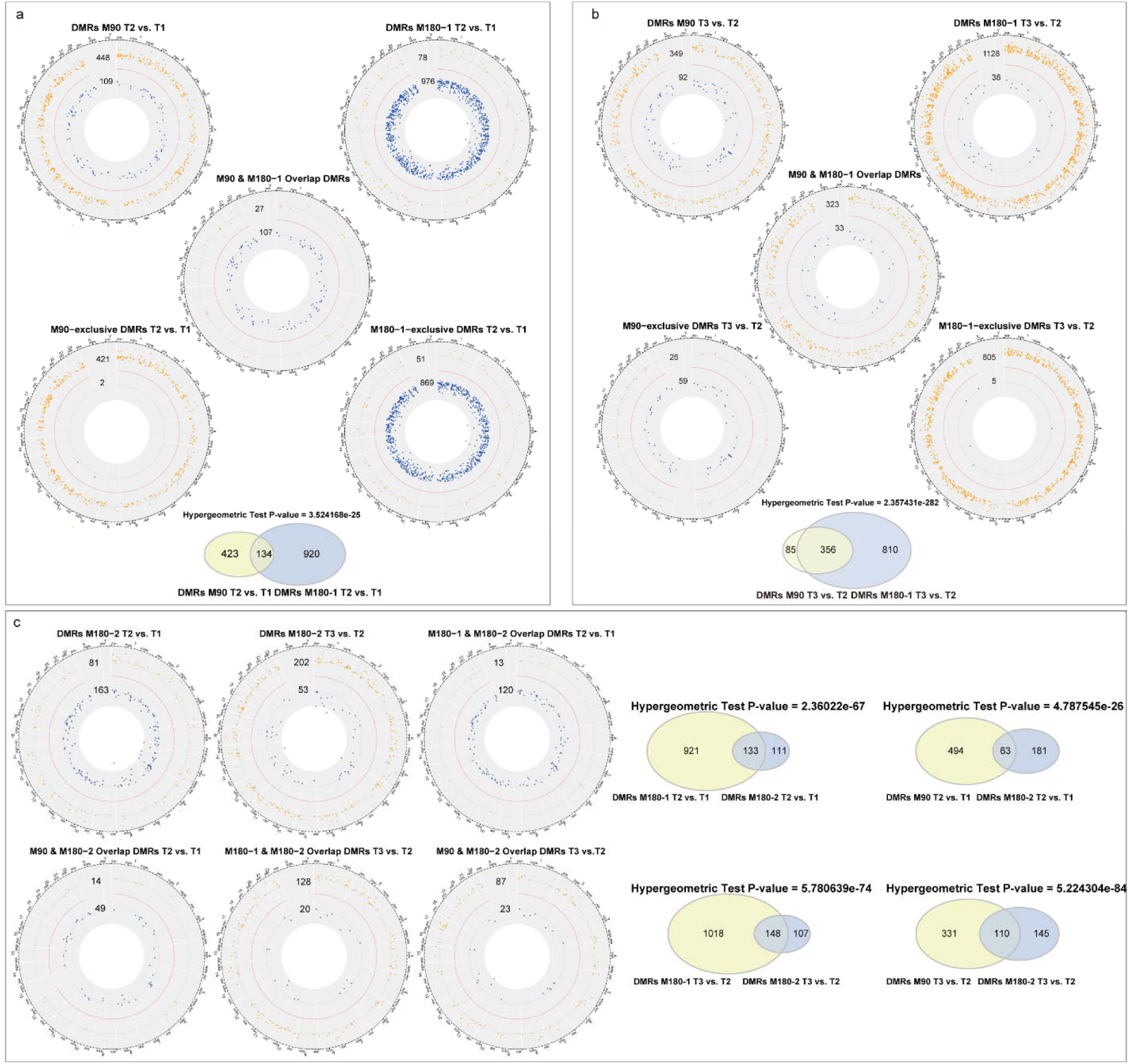
Genomic distributions of DMRs in M90, M180-1, and M180-2 **a,** Distribution of DMRs in the genome of M90 and M180-1 in T2 vs. T1. The y-axis represents the DMR values, with the red line indicating the zero-value threshold. The blue points correspond to hypomethylated DMRs, and the orange points represent hypermethylated DMRs. Distribution of DMRs present in both M90 and M180-1 (T2 vs. T1). This includes DMRs from M180-1 that overlap with those from M90 and DMRs from M90 that overlap with those from M180-1. Distribution of DMRs that are uniquely present in either the M90 or M180-1 T2 vs. T1. Number of overlapping and unique DMRs in M90 and M180-1 (T2 vs. T1). **b,** Distribution of DMRs in the genome of M90 and M180-1 in T3 vs. T2. **c,** Distribution of DMRs in the genome of M180-2 in T2 vs. T1 and T3 vs. T2, and overlap with DMRs in M90 and M180-1. The number of overlapping and unique DMRs between M180-2 and the other groups (M90, M180-1) is shown for both time comparisons.

**Figure S4.**
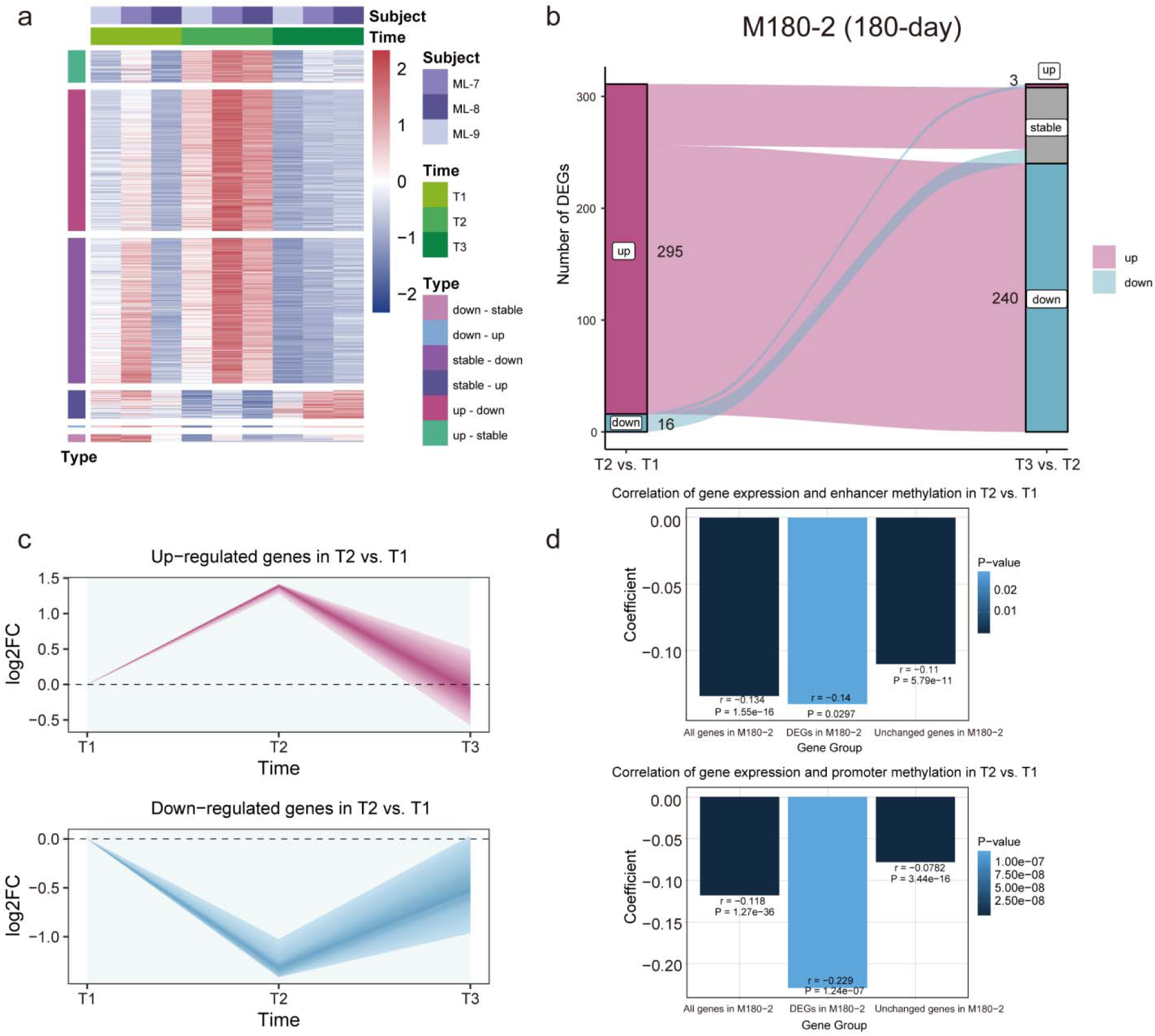
Alterations in gene expression and the correlation between DNA methylation and gene expression in an independent 180-day spaceflight (M180-2) **a,** The expression trends of all genes in spaceflight M180-2. Genes were grouped by the type of change in the T2 vs. T1 and T3 vs. T2 comparisons, e.g., “down_up” indicates downregulated in T2 vs. T1 and upregulated in T3 vs. T2, whereas “stable” indicates no significant change. **b,** Number of DEGs in T2 vs. T1 and the changes in these DEGs after recovery (T3 vs. T2) in M180-2. **c,** Dynamic changes in the expression levels of DEGs from T2 vs. T1 postflight (T2) and recovery (T3) in M180-2. **d,** The correlation coefficient of log_2_FCs of gene expression and log_2_FCs of promoter and enhancer area methylation in T2 vs. T1 was calculated for all expressed genes, all DEGs, and other unchanged genes.

**Figure S5.**
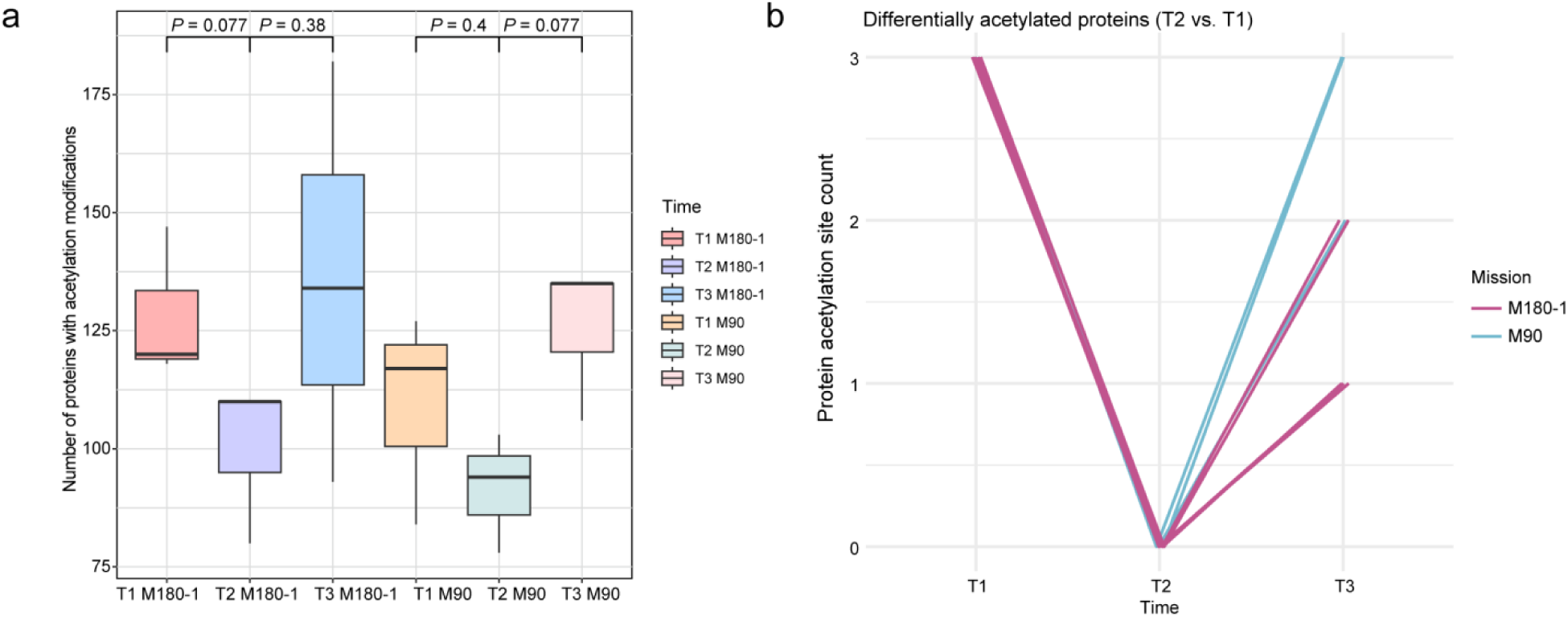
Changes in protein acetylation modifications in M90 and M180-1 **a,** The number of proteins with acetylation modifications in samples from the M90 and M180-1 spaceflights. **b,** Differentially acetylated proteins in M90 and M180-1. Each line represents a protein and the vertical axis corresponds to the number of acetylation sites of that protein. The source data for Figure S5 are presented in Table S5.

**Figure S6.**
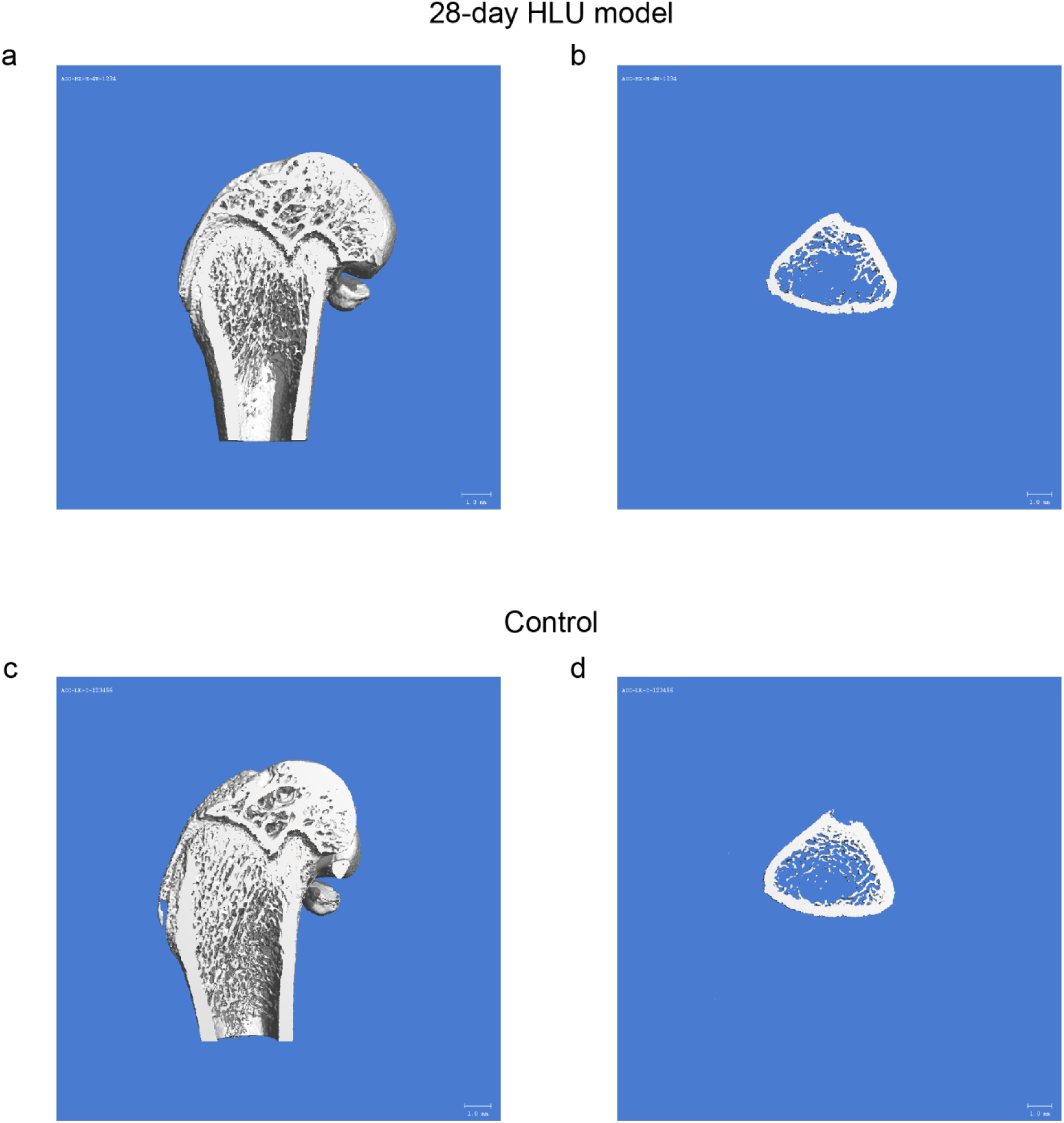
Reconstructed microCT images of bone morphology in 28-day HLU model rats and control rats Representative reconstructed microCT images of the longitudinal section of distal femurs collected from the groups of 28-day HLU model rats (**a**) and control rats (**c**). Images depicted cross sections of the same femurs from the HLU rats (**b**) and control rats (**d**), respectively.

**Figure S7.**
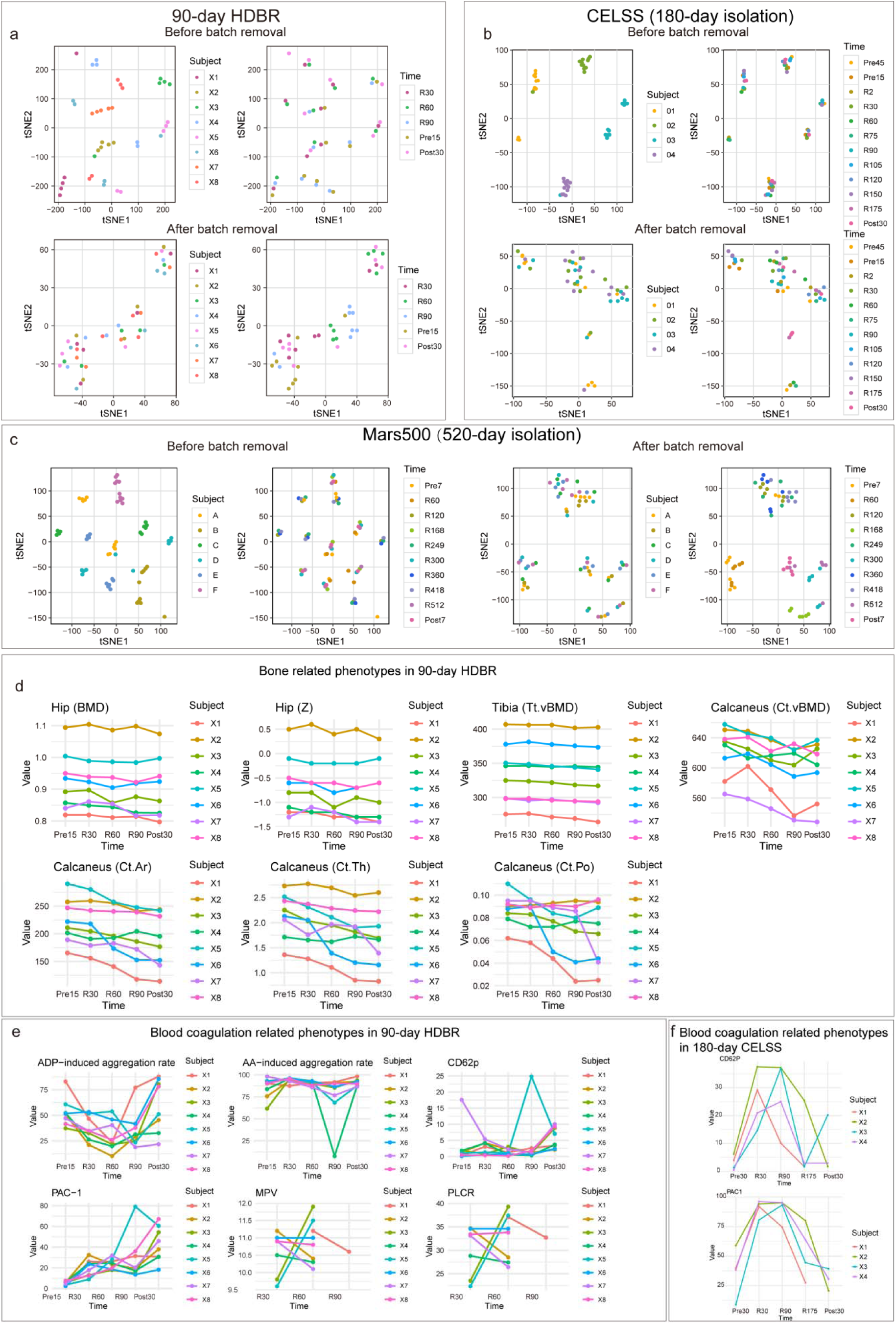
Global alterations in methylation and phenotype alterations in ground-based simulation experiments **a-c**, Demonstration of dimensionality reduction by tSNE before and after removal of batch effects for the 90-day HDBR (**a**), 180-day CELSS (**b**) and Mars500 (**c**). Annotations show the sample’s source (subject) and the time of sampling (time). **d-e**, Bone and coagulation related phenotype alterations in the 90-day HDBR. (**d**), Bone related phenotypes includes Bone Mineral Density (BMD) and Z-score of hip bone, tibia, and calcaneus. The prefix “Pre” of the measurement timepoint represents how many days before HDBR, “R” represents the number of days the experiment is conducted, and “Post” represents how many days after HDBR. (**e**), Coagulation related phenotypes includes ADP-induced platelet aggregation rate, AA-induced platelet aggregation rate, PAC-1 expression rate, CD62P expression rate, TAT Complex, MPV, and PLCR. (**f**), Coagulation related phenotype alterations in the 180-day CELSS. Coagulation related phenotypes includes PAC-1 expression rate and CD62P expression rate. The prefix “Pre” of the measurement time point represents how many days before the experiment, “R” represents the number of days the experiment is conducted, and “Post” represents how many days after the experiment.

## STAR⍰METHODS

### KEY RESOURCES TABLE

| REAGENT OR RESOURCE | SOURCE | IDENTIFIER |
| --- | --- | --- |
| Antibodies |  |  |
| Bacterial and virus strains |  |  |
| Biological samples |  |  |
| Chemicals, peptides, and recombinant proteins |  |  |
| Critical commercial assays |  |  |
| Deposited data |  |  |
| Transcriptomics data | This paper | HRA: HRA008486, HRA008487, HRA008488 (https://ngdc.cncb.ac.cn/gsa-human); OMIX: OMIX007304 (https://ngdc.cncb.ac.cn/omix); CRA: CRA018849, CRA018831 (https://ngdc.cncb.ac.cn/gsa) |
| DNA methylation data | This paper | OMIX: OMIX007289,<br>OMIX007291,<br>OMIX007300,<br>OMIX007301,<br>OMIX007290,<br>OMIX008613,<br>OMIX008614,<br>OMIX008615,<br>OMIX007302<br>(https://ngdc.cncb.ac.cn/omix) |
| Proteomics data | This paper | OMIX: OMIX007327,<br>OMIX007328<br>(https://ngdc.cncb.ac.cn/omix) |
| Protein acetylation data | This paper | OMIX: OMIX007319,<br>OMIX007320<br>(https://ngdc.cncb.ac.cn/omix) |
| Phenotypic data | This paper | OMIX: OMIX007309,<br>OMIX007310,<br>OMIX007312,<br>OMIX007306,<br>OMIX007307,<br>OMIX007359,<br>OMIX010025,<br>OMIX010026,<br>OMIX010027,<br>OMIX007311<br>(https://ngdc.cncb.ac.cn/omix) |
| Experimental models: Cell lines |  |  |
| Experimental models: Organisms/strains |  |  |
| Oligonucleotides |  |  |
| Recombinant DNA |  |  |
| Software and algorithms |  |  |
| R 4.2.1 | http://www.r-project.org/ | RRID:SCR_001905 |
| edgeR | http://bioconductor.org/packages/edgeR/ | RRID:SCR_012802 |
| UpSetR | https://github.com/hms-dbmi/UpSetR | RRID:SCR_026112 |
| ChAMP | http://www.bioconductor.org/packages/2.13/bioc/html/ChAMP.html | RRID:SCR_012891 |
| SVA | http://www.omicsexpress.com/sva.php | RRID:SCR_002155 |
| Nextflow | https://github.com/nextflow-io/nextflow | RRID:SCR_024135 |
| FastQC | http://www.bioinformatics.braham.ac.uk/projects/fastqc/ | RRID:SCR_014583 |
| MultiQC | http://multiqc.info/ | RRID:SCR_014982 |
| Trim Galore | http://www.bioinformatics.braham.ac.uk/projects/trim_galore/ | RRID:SCR_011847 |
| STAR | http://code.google.com/p/rna-star/ | RRID:SCR_004463 |
| RSEM | http://deweylab.biostat.wisc.edu/rsem/ | RRID:SCR_000262 |
| clusterProfiler | http://bioconductor.org/packages/release/bioc/html/clusterProfiler.html | RRID:SCR_016884 |
| WGCNA | http://www.genetics.ucla.edu/labs/horvath/CoexpressionNetwork/ | RRID:SCR_003302 |
| Cytoscape | http://cytoscape.org | RRID:SCR_003032 |
| Other |  |  |

### EXPERIMENTAL MODEL AND STUDY PARTICIPANT DETAILS METHOD DETAILS

#### Overview of the experiments and datasets

Here, we provide independent multi-omics data from multiple Chinese astronauts in one 13-day (M13), one 15-day (M15), one 33-day (M33), one 90-day (M90) and two 180-day spaceflights (M180-1 and M180-2) (Figure S1). For all the six spaceflights, DNA methylation and phenotypes were analyzed. For M90, M180-1 and M180-2, both transcriptome and DNA methylation were analyzed using peripheral blood cells, and the corresponding phenotypes were measured as well. For M180-1, proteome and protein acetylation were analyzed. Concurrently, we introduced and performed a variety of ground-based simulation experiments to estimate the effects of different space environment factors independently (Figure 1). The Mars500 (520-day isolation simulation^14^) and 180-day Controlled Ecological Life Support System (CELSS) ^13^ experiments were introduced to stimulate isolation in space. A 90-day head-down bed rest (HDBR) experiment and 28-day (rat) and 35-day (rat) hind-limb unloading (HLU) model experiments were conducted to simulate microgravity on Earth. Details of the ground-based simulation experiments are described in the Supplementary Methods.

All animal experimental protocols in this study were performed in accordance with standard ethical guidelines and approved by the China Astronaut Research and Training Center Animal Ethics Review Committee (NO: ACC-IACUC-2021-015).

#### Participants

A total of seventeen Chinese astronauts participating in one 13-day (M13, three astronauts), one 15-day (M15, three astronauts), one 33-day (M33, two astronauts), one 90-day (M90, three astronauts) spaceflight and two 180-day spaceflights (M180-1, three astronauts, and M180-2, three astronauts) were included. We calculated the statistics of the ages of all the seventeen astronauts from the six spaceflights. The minimum value is 33.42 years, the 1st quarter value is 41.75 years, the median value is 45.67 years, the mean value is 45.17 years, the 3rd quarter value is 48.75 years, the maximum value is 56.75 years, the standard deviation is 6.73 years. There exists no outlier value among all the seventeen age values, indicating consistent age range. The study subjects signed informed consent according to the declaration of Helsinki and Article 1008 Civil Code of the People’s Republic of China for the collection and use of sample materials and data in research protocols at China Astronaut Research and Training Center and the collaborating institutions. The study protocols were approved by the China Astronaut Research and Training Center Animal Medical Ethics Review Committee.

#### Sample collection

For each of the six spaceflights, blood samples from each astronaut at three or four timepoints (Table S1) were collected into 2 mL BD Vacutainer® EDTA Tubes (Cat # 367841) per the manufacturer’s recommendations. For the latest three spaceflights (M90, M180-1 and M180-2), additional blood samples were collected into 2.5 mL BD PAXgene® Blood RNA Tubes (Cat # 762165) for transcriptome sequencing. Sample collection was performed at 6:30 o’clock a.m. each time to avoid circadian rhythmic effects. To minimize batch effects within each spaceflight, the blood samples were stored at −80 ℃ until all the samples from the same spaceflight were fully prepared.

#### Multi-omics measurement

For multi-omics data acquisition, DNA methylation was profiled via the Illumina Infinium MethylationEPIC BeadChip. Samples from the earlier spaceflight missions (M13, M15 and M33) were analyzed using 450K BeadChip, whereas, later spaceflight missions (M90, M180-1 and M180-2) were analyzed using the 850K BeadChip, reflecting the updates of the DNA methylation assay by Illumina. RNA sequencing was conducted on an Illumina NovaSeq 6000 PE150 platform using the standard method. Proteomics and protein acetylation were quantified on a Bruker timsTOF Pro mass spectrometer. The full details are available in the Supplementary Information.

## QUANTIFICATION AND STATISTICAL ANALYSIS

### Expression quantification and differential expression analysis

To facilitate transcriptomic analysis, the following systematic pipeline was employed for the preprocessing of raw RNA sequencing data, utilizing the human genome from Ensembl version 107 as the reference.

FastQC software (v0.12.1) (https://www.bioinformatics.babraham.ac.uk/projects/fastqc/) was used to evaluate the quality of the original FASTQ files. Using parameters including --length 36, a quality threshold of --quality 25, and a trimming stringency of --stringency 3, TrimGalore software (v0.6.10) (https://github.com/FelixKrueger/TrimGalore/releases/tag/0.6.10) was used to trim and filter the initial FASTQ files, efficiently eliminating low-quality nucleotides and adapter sequences. STAR (v2.7.10a) (https://github.com/alexdobin/STAR/)^56^ was modified to align the trimmed FASTQ files to the Ensembl version 107 reference genome (hg38). The RSEM tool (v1.3.1) (https://github.com/deweylab/RSEM)^57^ was used to compute gene expression abundance according to matched BAM files to generate an expression profile. The above analysis was performed on CentOS Linux version 7.2.1511. The Rtsne package (v0.16) was used for dimensionality reduction and visualization. To mitigate the potential influence of interindividual variability and batch effects, the combat method was employed. We used ComBat_seq from the sva package (v3.46.0)^58^ to remove the individual-specific characteristics from each dataset by treating individual sources as batches to remove these environmentally irrelevant effects. After batch effect removal, the two flight datasets were merged for further analysis. Differential expression analysis was conducted using the limma package (v3.54.2)^59^, and DEGs (|log_2_FC| > log2(1.5), and adjusted *P* value < 0.05, DEGs) were identified across all groups. All of the R packages used in this study were applied in R version 4.2.1 (https://cran.r-project.org/bin/windows/base/old/4.2.1/). The DEG classification method is available in the Supplementary Methods.

### Alternative splicing analysis

The clean RNA-seq data obtained via the above TrimGalore processing were subjected to alternative splicing analysis using the TopHat Cufflinks pipeline^60^. The analysis was carried out with Bowtie2-2.5.1-linux-x86_64 (https://sourceforge.net/projects/bowtie-bio/files/bowtie2/2.5.1/), TopHat v2.1.1 (https://ccb.jhu.edu/software/tophat/downloads/), and Cufflinks v2.2.1 (https://cole-trapnell-lab.github.io/cufflinks/install/) on CentOS Linux release 7.2.1511. After quality control, the clean FASTQ data were first aligned using TopHat to the reference genome Homo_sapiens.GRCh38.dna_sm.primary_assembly.fa, which was downloaded from ENSEMBL, with guidance from Homo_sapiens.GRCh38.107.gtf, which contains transcript information. The Cufflinks package was subsequently used to assemble transcripts, compare the assemblies to annotations, merge the assemblies and detect differential splicing and promoter use. The differentially expressed alternatively spliced transcripts were counted and then enriched with STRING (https://cn.string-db.org/).

### DNA methylation raw data processing

DNA methylation data from space- and human ground-based experiments were processed with ChAMP and then visualized with the Rtsne package (0.16) in R version 4.2.1^61,62^ for dimensionality reduction. As in the transcriptomic data, there was a batch effect between the two flight datasets, with samples in each flight grouped primarily according to individual sources. To remove the batch effects, we carried out a procedure similar to that used for the transcriptome data, applying the champ.runCombat method in the ChAMP package (2.28.0) (R version 4.2.1).

### Differential methylation analysis

Differential methylation analysis was performed using ChAMP. Positions with *P* value < 0.05, |deltaBeta| > 0.05, and a consistent change across all three subjects (either all demethylated or all hypermethylated) were considered differentially methylated between the two groups and defined as differentially methylated positions (DMPs). Differentially methylated regions (DMRs) were identified using the champ.DMR function, employing the Bumphunter method with an adjusted *P* value threshold set to 0.05. DMRs with containment relationships (including complete overlap) in the two flight experiments were identified by comparing their positions on the chromosome. The functional enrichment of DMRs was performed on the genes hosting the DMPs distributed in the region via the clusterProfiler package (v4.6.2)^63^.

### Methylation change trend analysis

We compared the trends in genomic methylation changes in spaceflight (M90 and M180-1) and ground-based simulation experiments (90-day HDBR, 180-day CELSS, and Mars500) for a set of genes enriched by nodes of the adaptation-related PPI network, that is, the functional genes of interest. To select appropriate loci exhibiting consistent changes, the following criteria were employed: loci on the promoter region (including “TSS1500”, “TSS200”, “5’UTR”, and “1stExon”) with a corrected *P* value < 0.05 as determined by an ANOVA test and grouped by time, as well as loci exhibiting a range of fluctuating beta values greater than 0.05. The DNA methylation levels of these selected loci were subsequently averaged across a group of experimental subjects to facilitate visualization of the data. The visualization method began with data normalization of the matrix of average methylation values for the loci, followed by the construction of a graph depicting the trends in the methylation levels of the locus over time. The graphs used filled regions from light to dark to represent the range of intervals of loci methylation levels, illustrating changes in gene set methylation levels over time.

### Correlation between DNA methylation and expression

The significance of the connection between promoter region methylation and expression was evaluated by performing a Pearson correlation test on the log_2_FC values of gene expression and the log_2_FC values (deltaBeta) of promoter/enhancer region DNA methylation (the median deltaBeta of loci on the promoter/enhancer region). DMPs located in the promoter regions were automatically annotated by the champ.DMP function. We identified DMPs in the enhancer regions through annotating against the human-specific FANTOM5 enhancer database^64^.

### Proteomics and protein acetylation data processing

The data from the machine were identified using MaxQuant’s (v.1.5.3.30) integrated Andromeda engine; then, quantitative analysis was performed using MaxQuant on the basis of the peak intensity, peak area and retention time of peptide segments related to primary mass spectrometry, and the targeted modified peptide segments were extracted. We used the UniProt protein database and genome annotation-based protein database for protein identification. The data obtained were filtered using the Andromeda engine integrated in MaxQuant, filtering at the PSM-level with an FDR <= 1% at the spectrum level and further filtering at the protein-level FDR <= 1% at the protein level. Additionally, for modification sites, filtering was performed at the modification site level with an FDR of 1% for identification (site decoy fraction <= 1%) to obtain significant modification results. The modified peptides were counted for further differential analysis.

### Differential expression analysis of proteomics data

The differential expression analysis of expressed peptides was performed with limma (v3.54.2) in R version 4.2.1. Quantile normalization was first applied to the data before linear regression. The significance threshold for differentially expressed peptides was an adjusted *P* value <= 0.05.

### Differential expression analysis of protein acetylation data

The differential fold changes in the modified peptide segments in each comparison group were subsequently calculated on the basis of the defined comparison groups and statistically tested using edgeR 3.40.2. The data were first quantile normalized with the voom function, and then a linear model was fitted to the expression data for each gene. An adjusted *P* value of less than 0.05 was used as the threshold for differentially modified peptides.

### Construction of PPI networks related to environmental adaptation

Initially, the WGCNA package (v1.72-5)^65^ was employed to partition genes into modules consisting of closely associated genes. Subsequently, the correlation *P* values between all modules and time (T1, T2, T3) were calculated. The brown and blue modules were identified on the basis of a *P* value threshold for significance of 0.05. The ClusterProfiler (v4.6.2)^63^ package was used to perform functional enrichment analysis of the module genes. The simplifyGO function from the simplifyEnrichment^66^ (v1.8.0) package was employed to cluster and visualize the GO terms. The biological functions within each cluster were summarized and presented as word clouds, which were then linked to a similarity heatmap displaying the shared biological functions across the clusters. The importance of each keyword was determined using Fisher’s exact test, and their significance was represented by the size of the font used.

To investigate the interplay between different functional genes within the brown module, gene-gene interactions were assessed by integrating the gene coexpression coefficients, which were calculated using WGCNA, with the gene interaction scores from the STRING database. A threshold of WGCNA weights > 0.125 (equivalent to the absolute value of the coexpression coefficient > 0.5 where power = 3) and a STRING score > 0.4 were applied to identify interacting genes, among which 1727 genes were identified. This step ultimately led to the construction of a predicted protein-protein interaction (PPI) network, with a total of 12490 interactions and 1458 nodes.

### Gene set interaction analysis

To calculate the interaction scores between geneset1 and geneset2, the following steps were taken. First, sub_PPI_net, a PPI network subset, was extracted containing genes from both geneset1 and geneset2. The calculation formula for the interaction score is as follows:

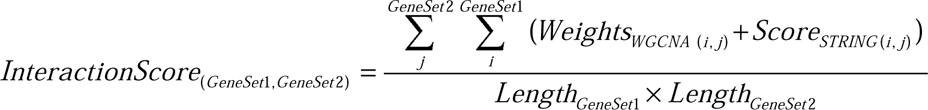

To assess the significance of the interaction score, 100 random gene sets with the same size as geneset1 and geneset2 were randomly generated, and the interaction scores were calculated for these sets. The frequency of actual interaction scores greater than the scores obtained from the 100 random sets was calculated as the *P* value for interaction significance. Gene set pairs with an interaction score *P* less than 0.05 were considered to have significant interactions. The importance of each gene set was determined by the sum of the interaction scores between the gene set and all other gene sets.

