## Supplementary material for "Dynamic alterations of DNA methylation and transcriptome in adaptation to and recovery from the space environment": Supplemetary Information & Table S1

**Supplemental Information**

### Supplemental Methods

#### Details of multi-omics measurement for astronauts

**Nucleic acid extractions and quality control.** Total RNA was extracted from 2.5 mL blood in BD PAXgene® Blood RNA Tubes (Cat # 762165, BD, USA) using the TRIzol reagent (Invitrogen, CA, USA) according to the manufacturer’s protocol. RNA purity and quantification were evaluated using the NanoDrop 2000 spectrophotometer (Thermo Fisher Scientific, Waltham, MA, USA). RNA integrity was assessed using the Agilent 2100 Bioanalyzer (Agilent Technologies, Santa Clara, CA, USA).

Genomic DNA was extracted from 0.5 mL blood in BD Vacutainer® EDTA Tubes (Cat # 367841, BD, USA) using QIAamp DNA Blood Mini Kit (Qiagen, Germany) according to the manufacturer’s protocol. DNA concentration and integrity were assessed by a NanoDrop 2000 spectrophotometer (Thermo Fisher Scientific, Waltham, MA, USA) and agarose gel electrophoresis, respectively.

**Protein extraction and quality control.** Appropriate amounts of L3 lysis buffer (7 M urea, 2 M thiourea, 20 mM Tris–HCl, pH 8.0) without SDS were added to the 0.5 mL blood in BD Vacutainer® EDTA Tubes (Cat # 367841, BD, USA), followed by 2 mM ethylenediamine tetra acetic acid (Amersco, USA) and 1% protease inhibitor cocktail (Roche, USA), and incubated on ice for 5 min. Samples were ground with a grinder (60 Hz, 2 min) and centrifuged at 25000 g for 15 min at 4 °C. Next, 10 mM dithiothreitol (Amersco, USA) was added, and the mixture was reacted at 56 °C for 1 hour. Next, 55 mM iodoacetamide (Amersco, USA) was added and reacted for 45 min in the dark at room temperature (RT). Quantitative electrophoresis was then performed. The concentration of protein extract was measured using the Bradford assay^1^. The purity of the extracted proteins (10 μg) was verified and the proteins were separated by 12% SDS–PAGE, followed by Coomassie blue staining. For protein enzymatic digestion, 100 μg of protein per sample was digested with 1 μg of trypsin enzyme (protein/enzyme ration of 100:1) overnight at 37 °C. Finally, enzymatic peptides were desalted using a Strata X column and vacuum dried prior to mass spectrometry.

**DNA methylation beadChips.** Peripheral blood DNA of subjects in spaceflights were collected, 500 ng DNA of each sample was used to bisulfite converted using Zymo Research EZ DNA methylaiton-Glod Kits (Zymo Research, Irvine, CA, USA), then converted products were put into Illumina Methylation Arrays for measurements of DNA methylation. Finally, Illumina iSCAN (Illumina, USA) was used to scan the chip to get the Idat files.

**RNA sequencing.** RNA library for RNA-seq was prepared as rRNA depletion and stranded method. Briefly, the ribosomal RNA was depleted from total RNA using Illumina Ribo-Zero plus rRNA Depletion Kit (Illumina, USA) following manufacturer’s instruction. Library construction including RNA fragmentation, reverse transcription, adenylation of 3’ ends of DNA fragments, sequencing adaptor ligation and purification were conducted using Hieff NGS Ultima Dual-mode mRNA Library Prep Kit for Illumina (Yeasen, China) following manufacturer’s instruction. After library construction, the concentration of library was measured by the Qubit fluorometer (Qubit 4.0, Invitrogen, USA). The accurate concentration of cDNA library was again examined using qPCR. The size distribution of library was detected by agarose gel electrophoresis. After library preparation and pooling of different samples, the samples were subjected for Illumina sequencing. The RNA-seq were conducted on Illumina NovaSeq 6000 PE150 Platform (Illumina, USA).

**Blood proteomics.** *Liquid Chromatography.* All proteomic experiments were performed using a Bruker nanoElute (nanoElute 2, Bruker, Germany). The dried peptide samples were reconstituted with mobile phase A (100% H_2_O, 0.1% formic acid), centrifuged at 20000 g for 10 min, and the supernatant was taken for injection. The sample was first enriched in the trap column and desalted, and then entered a tandem self-packed C18 column, and separated at a flow rate of 300 nL/min by the following effective gradient: 0 min, 2% mobile phase B (100% acetonitrile, 0.1% formic acid); 0~45 min, mobile phase B linearly increased from 2% to 22%; 45~50 min, mobile phase B rose from 25% to 35%; 50~55 min, mobile phase B rose from 35% to 80%; 55~60 min, 80% mobile phase B. The nanoliter LC separation end was directly connected to the mass spectrometer and detected according to the following parameters:

*DIA Mass Spectrometry Detection.* The peptides separated by liquid phase chromatography were ionized by a nanoESI (Electrospray Ionization) source and then passed to a tandem mass spectrometer TimsTOF Pro (nanoElute 2, Bruker, Germany) for the Data Independent Acquisition (DIA) mode detection. The main parameters were set: ion source voltage was set to 1.6 kV, ion mobility range was 0.6-1.60 V.S/cm^2^, and MS1 mass spectrometer scanning range was 302~1077 m/z. The peak intensity above 2500 can be detected. The 302~1077 m/z was divided into 4 steps, and each step was divided into 8 Windows. A total of 32 Windows was used for continuous window fragmentation and information collection. The fragmentation mode was CID, the fragmentation energy was 10 eV, and the mass width of each window was 25. The cycle time of a DIA scan was 3.3 s. This process is based on the sample data generated from a high-resolution mass spectrometer.

**Blood acetylation modification quantification proteomics.** *Peptide enrichment.* Dissolve peptide segment with IAP buffer (IG125-10G, SIGMA, USA), ensure pH value is about 7. The peptide solution was mixed with microbeads (13416S-Acetyl-LysineMotif [Ac-K] Kit, Cell Signaling Technology, USA), incubated at 4 °C for 2 hours, centrifuge at 4 °C with 2000 g for 30 s, and discard the supernatant. The microbeads were cleaned twice with IAP buffer and three times with ice water. Add 55 μL 0.15% TFA into microbeads, incubate at RT for 10 min, and mix several times every 2 to 3 min. Centrifuge, collect supernatant, repeat once, merge supernatant. The eluted peptide solution is purified with ZipTip (Millipore, USA) and drained in a drain machine.

*Liquid Chromatography.* All proteomic experiments were performed using a Thermo UltiMate 3000 UHPLC (Thermo Fisher Scientific, USA). The dried peptide samples were reconstituted with mobile phase A (2% acetonitrile, 0.1% formic acid), centrifuged at 20000 g for 10 min, and the supernatant was taken for injection. The sample was first enriched in the trap column and desalted, and then entered a tandem self-packed C18 column. Then it was separated at a flow rate of 300 nL/min by the following effective gradient: 0~5 min, 5% mobile phase B (98% acetonitrile, 0.1% formic acid); 5~45 min, mobile phase B linearly increased from 5% to 25%; 45~50 min, mobile phase B rose from 25% to 35%; 50~52 min, mobile phase B rose from 35% to 80%; 52~54 min, 80% mobile phase B; 54~60 min, 5% mobile phase B. The nanoliter liquid phase separation end was directly connected to the mass spectrometer and detected according to the following parameters:

*Mass Spectrometry Detection.* The peptides separated by liquid phase chromatography were ionized by a nanoESI source and then passed to a tandem mass spectrometer TimsTOF Pro (nanoElute 2, Bruker, Germany) for DDA (Data Dependent Acquisition) mode detection. The main parameters were set: ion source voltage was set to 1.6 kV, MS1 mass spectrometer scanning range was 100~1700 m/z; ion mobility range was 0.6-1.60 V.S/cm^2^; MS2 mass spectrometer scanning range was 100~1700 m/z; The precursor ion screening conditions for MS2 fragmentation: charge 0 to 5+, and the first 10 precursor ions with the peak intensity exceeding 10000 and the peak intensity above 2500 can be detected. The MS1 cumulative scan time was 100 ms; MS2 cumulative scan time was 100 ms; The ion fragmentation mode was CID (Collision-Induced Dissociation), and the fragment ions were detected in TOF. The dynamic exclusion time was set to 30 s.

**Phenotypes.** “Phenotypes” monitored for M13, M15, M33, M90, M180-1 and M180-2 astronauts encompass a wide range of biochemical markers, physiological parameters, and functional assessments, which can be broadly categorized into several groups: hormonal and endocrine factors, bone and mineral metabolism indicators, cardiovascular markers, muscular and skeletal parameters, and biochemistry, metabolic profiles. Specifically, hormonal and endocrine factors such as adrenocorticotropic hormone, antidiuretic hormone, and cortisol; bone metabolism indicators like alkaline phosphatase and osteocalcin; cardiovascular markers such as angiotensin II, blood pressure, and heart rate variability; muscular and skeletal parameters including muscle circumference and bone density; blood biochemistry like white blood cell interleukins, hemoglobin, cellular adhesion factors, growth factors, antioxidant capacity; metabolic profiles encompassing glucose, cholesterol, and triglycerides. Overall, these diverse parameters provide a comprehensive evaluation of various physiological systems states of astronauts throughout spaceflight (Links of downloadable phenotype indicators and datasets are listed in https://www.spacelifescience.cn/datasets).

**Human bone phenotypic measurement.** Bone densities of lumbar spine and hip were measured by Dual Energy X-ray Absorptiometry (DXA, Osteocore-1, Medilink, France). The scan of lumbar spine and hip was performed according to standard scanning methods, ROIs were automatically selected and bone density results were analyzed by the DXA built-in program. Calcaneal bone density was measured by ultrasound bone sonometers (Pegasus Smart, Medilink, France) according to standard scanning method.

**ELISA analysis.** For human thrombin-antithrombin complex (TAT) ELISA analysis, serum was collected and stored at –80 °C until use. The ELISA was performed by a conventional colorimetric detection using Human TAT ELISA kit (Keqiaobio, China) according to the manufacturer’s instructions. In brief, 50 µl of serum were pipetted in duplicate into the wells of the precoated ELISA plate. Then, 50 µl of antibody solution were added to each well and incubated at RT for 90 min on a shaking device. After incubation, the plates were washed three times with PBS. Finally, the absorbance of each well was read by Multiskan Spectrum (Thermo Fisher Scientific, USA).

**Human CD62P detection.** Fresh blood collected by Vacutainer Citrate Tube (363095, BD, USA) from spaceflights was added into a FACS tube containing PerCP labeled mouse anti-human CD61 antibody (340506, BD, USA) and PE labeled mouse anti-human CD62P antibody (348107, BD, USA) and incubated at RT for 20 min in the dark. Then, blood was fixed with 1% cold paraformaldehyde and analyzed by flow cytometry (FACS Aria Ⅱ, BD, USA). As negative control, fresh blood was incubated with PerCP labeled mouse anti-human CD61 antibody (340506, BD, USA), PE labeled mouse anti-human IgG antibody (340013, BD, USA).

#### Materials from 90-day head-down bed rest (HDBR)

**Participants.** A group of eight male volunteers with education levels beyond high school were selected for a 90-day bed rest experiment conducted at -6° head-down tilt (HDT). Prior to their participation, these volunteers underwent a thorough physical examination, including routine medical and laboratory tests, to ensure they did not have chronic diseases such as neurological disorders, musculoskeletal system disorders, infectious diseases, or dyssomnia. Furthermore, they were all non-users of medications, drugs, tobacco, and alcohol.

A comprehensive explanation of the study, including its research purpose, experimental procedures, methods, conditions, potential problems, and complications, was provided to the participants. Each volunteer subsequently provided written informed consent. Upon completion of the study, the participants received financial compensation. The study protocol adhered to the ethical guidelines outlined in the 1975 Declaration of Helsinki and was approved by the Institutional Review Board of the China Astronaut Research and Training Center.

**Blood** **collection and measurement.** After collecting peripheral blood samples into EDTA-containing tubes, the tubes were subjected to centrifugation at 1200 g for 15 min at RT within 30 min of collection. The resulting supernatant fluid, which contained plasma, was carefully transferred into microcentrifuge tubes. To eliminate cellular debris, a second centrifugation was performed at 12000 g for 10 min at 4 °C. The plasma was subsequently divided into aliquots and stored at -80 °C until required for use. DNA methylation was profiled using Illumina Infinium MethylationEPIC BeadChip (see details in “DNA methylation beadChips” from “Details of multi-omics measurement for astronauts”).

#### Materials from 180-day Controlled Ecological Life Support System Experiment (CELSS)

**Subjects and experiments.** From June to December 2016, a 180-day CELSS Experiment was conducted at the SPACEnter Space Science and Technology Institute in Shenzhen, China, involving four subjects (three males and one female) aged 29–43 years. During the experiment, the participants experienced 180 days of isolation and adhered to a Mars solar day schedule from 10 : 30 PM on day 71 to 10 : 30 PM on day 108. The CELSS consisted of six cabins, including one for the crew and life support system for daily activities, cooking, and sleeping. Another cabin was dedicated to the resource system, utilized for processing nonedible plant materials and generating CO_2_. Additionally, four greenhouses were utilized to cultivate 25 different plant varieties. The indoor air in each cabin was typically not ventilated, except when the oxygen ratio fell below 19.0% or the CO_2_ level dropped below 500 ppm. For further details regarding the subjects and experimental environment, refer to previous descriptions^2-4^. Throughout the study, the four subjects abstained from smoking and consuming alcohol.

**Sample collection and measurement.** Serum samples were collected from the four fasting subjects at various timepoints: 30 days before the mission (-30d), 2d, 30d, 60d, 75d, 90d, 105d, 120d, 150d, and 175d during the mission, and 30 days after the mission (+30d). Additionally, strict control measures were implemented for factors such as food intake, physical activity, temperature, and the sleep-wake cycle prior to each blood collection. To ensure proper storage, all collected samples were kept at -80 °C until further analysis. DNA methylation was profiled using Infinium Methylation 450K assay (see details in “DNA methylation beadChips” from “Details of multi-omics measurement for astronauts”).

Transcriptome expression was quantified using Affymetrix Human Transcriptome Array 2.0 (HTA2.0). The fragmented and biotinylated RNA samples were prepared according to the standard Affymetrix GeneChip WT PLUS Reagent Kit (Affymetrix, USA) protocol from 100 ng total RNA starting material (Mars500 samples) and 5.5 μg cDNA intermediate product. DNA targets were hybridized for 17 hours at 45 °C on GeneChip Human Transcriptome Arrays 2.0 (Affymetrix, USA). GeneChips were washed and stained in the Affymetrix Fluidics Station 450 (Affymetrix, USA) according to the standard GeneChip Expression Wash, Stain and Scan Kit (Affymetrix, USA) protocol. Subsequently, the GeneChips were scanned using the Affymetrix 3000 7 G scanner (Affymetrix, USA). Raw CEL files were further analyzed.

#### Materials from the Mars500 project

**Mission background.** The Mars500 project^5^, conducted by the Institute of Biomedical Problems (IBMP) of the Russian Academy of Sciences (RAS) in Moscow, simulated an interplanetary manned flight with a focus on the crew members’ health and working capacity. Data collected during the three stages of the simulated flight (flight to, landing at, and return from Mars) were analyzed. The six-member crew consisted of three Russians, one Italian, one French, and one Chinese individual. All scientific experiments were approved by the IBMP committee on Bioethics, and the crew members underwent clinical examinations and signed consent forms before participating.

The simulation took place at the IBMP isolation facility in Moscow, Russia. Various periods were simulated throughout the 520-day isolation, including flight along a spiral path in the Earth’s gravitational field (1d–11d), flight towards Mars on a heliocentric trajectory (51d–204d), flight along a spiral path in the Martian gravitational field (205d–243d), flight in Mars’ orbit, including simulated trips to the Martian surface by two crew members (244d–272d), flight along a spiral path in the Martian gravitational field (273d–309d), flight towards Earth on a heliocentric trajectory (310d–467d), and flight along a spiral path in the Earth’s gravitational field (468d–520d).

**Biochemical and epigenetic assay.** Blood samples were collected from all six crew members at multiple time points during the experiment. The collection days were -7, +60 (+61), +120 (+121), +168 (+169), +249 (+250), +300 (+301), +360 (+361), +418 (+419), +510 (+511), and +527. The crew members were randomly divided into two groups on consecutive days for each blood extraction. After extraction, the blood cells were treated with EDTA anticoagulation, centrifuged at 4 °C, and the plasma was separated.

For DNA extraction, frozen blood samples were treated with proteinase K/RNase and phenol/chloroform. Bisulfite modification was conducted using an EZ DNA Methylation Kit, and the Infinium Methylation 450K assay was performed according to Illumina’s protocol.

#### Multi-omics measurement for HLU model

**HLU model.** Male SD rats aged 7 weeks were allowed to acclimatize their new surrounding for 1 week as singletons and free access to water and standard chow under a 12/12 hours light/dark cycle. Subsequently, the rats were randomly assigned to the control group (CN) and the HU1 group and HU2 group. The experimental manipulation followed the methodology of Wronski and Morey-Holton. Briefly, the rat was suspended by the tails using a strip of adhesive surgical tape attached to a chain hanging from a beam. The rats were 30° angle to the floor with only the forelimbs touching the floor and allowed to move freely to food and water. HU1 were suspended first, HU2 were suspended 7 days after. After 28-day suspension for HU2 and 35-day suspension for HU1, all rats were euthanized for blood collection and femur bone dissection.

**Sample collection and measurement.** At the end of the experiments, rat’s blood was collected from heart for sequencing into 2 mL BD Vacutainer® EDTA Tubes (Cat # 367841) and 2.5 mL BD PAXgene® Blood RNA Tubes (Cat # 762165) per manufacturer’s recommendations. RNA sequencing was conducted on Illumina NovaSeq 6000 PE150 Platform for all rats (see details in “RNA sequencing” from “Details of multi-omics measurement for astronauts”). For HU2 group and CN group, bone morphological parameters of femur were obtained by μCT system vivaCT40 (SCANCO medical AG, Switzerland). Blood routine and coagulation factors in HU2 group and CN group were performed by hematology analyzer. The concentration of P-selectin, TXB2 and PF4 were detected in blood serum by ELISA kits (using Rat P-selectin ELISA Kit (Ruixinbio, China), Rat TXB2 ELISA Kit (Ruixinbio, China) and Rat PF4 ELISA kit (Ruixinbio, China), see details in “ELISA analysis” from “Details of multi-omics measurement for astronauts”).

**Rat femur μCT analysis.** The 28-day HLU model rat femur bone was scanned ex vivo using an μCT system vivaCT40 (SCANCO medical AG, Switzerland) with an isotropic voxel size of 10.5 μM. The 3D reconstruction of the mineralized tissue was performed automatically by the system. About 160 slices of distal femora metaphysis was chosen for analysis of the microarchitecture parameters, including Bone Mineral Density (BMD), Bone Volume Fraction or Bone Volume/Total Volume (BV/TV), Trabecular Number (Tb.N) and Trabecular Separation (Tb.Sp).

#### Classification of DEGs

We classified all expressed genes into DEGs (differentially expressed at T2 vs. T1 or T3 vs. T2) and unchanged genes, and DEGs were categorized as unrecovered DEGs, over-range rebound DEGs and other DEGs based on the alteration of DEGs in both T2 vs. T1 and T3 vs. T2 phases. A number of genes were changed but didn’t reach statistical significance at T2 vs T1, but were significantly differentially expressed at T3 vs. T2 due to overrange rebound. Thus, overrange rebound genes were characterized as genes that were downregulated at T3 (compared with T2) and had a lower expression level than T1, as well as genes that were upregulated at T3 (compared with T2) and had a greater expression level than T1. Unrecovered genes were classified as those that were significantly changed at T2 vs. T1 but not significantly differentially expressed at T3 vs. T2. Other DEGs were considered to be mostly recovered and not exceeding initial levels in T1.

Raw RNA sequencing data from rats, using the Ensembl version 107 of the rat genome as a reference, were processed in the same procedure as above. Differential expression was also calculated by the limma package.

### Supplemental Notes

#### Note S1. A widespread decrease in DNA methylation levels after spaceflight

**Statistics of probe**’**s** **genomic functional regions****.** For differentially methylated probes, the proportion of the probes that are located in the promoter region (including "TSS1500", "TSS200", "5’ UTR", and "1stExon") was counted as a standalone subject (Figure S1b). When the deltaBeta threshold was set to 0.05, it was found that T2 vs. T1 hypomethylated loci (M90, 25.0%; M180-1, 27.2%, respectively) distributed in the promoter regions more frequently than hypermethylated loci (M90 20.4%; M180-1 18.6%, respectively) did in both flights, whereas T3 vs. T2 hypermethylated loci (M90, 28.2%; M180-1, 29.6%, respectively) distributed in the promoter regions more frequently than hypomethylated loci (M90, 19.2%; M180-1, 17.6%, respectively). This observation held consistent with hypo/hypermethylated loci in the enhancer regions (Figure S1b). Similarly, it was found that T2 vs. T1 hypomethylated loci (M90, 14.2%; M180-1, 11.1%, respectively) distributed in the enhancer regions more frequently than hypermethylated loci (M90, 7.4%; M180-1, 6.5%, respectively) in both flights, whereas T3 vs. T2 hypermethylated loci (M90, 12.2%; M180-1, 10.7%, respectively) distributed in the enhancer regions more frequently than hypomethylated loci (M90, 5.7%; M180-1, 5.7%, respectively).

**Dysregulation of expression of methylation-related enzymes in M180-2 and HLU model rats.** *TET2* (log_2_FC=0.604, adjusted *P* value =0.002, limma) and *TET3* (log_2_FC=0.461, adjusted *P* value =0.005, limma) were significantly increased (T2 vs. T1) in M180-2 (Table S3). *TET2* (Tet2 in rat) expression was also significantly increased in 35-day HLU model rats compared to the control group (log_2_FC=0.256, *P*=0.025, limma) (Table S7). Another DNA Methyltransferases *DNMT3A* (Dnmt3a in rat) was significantly decreased in 28-day HLU model rats (log_2_FC=-0.749, *P*=4.402e-4, limma) (Table S7). As a result, the simulated microgravity has the potential to affect methylation enzymes.

#### Note S2. Mathematical modeling for overrange rebound

Overshoot is a general concept in ecology, economics, engineering, and generally refers to the extent to which a target goal or limit is exceeded. To quantify the overshoot for each gene, we used the following equation:

$Overshoot=Maximum E_{overshoot}/E_{amplitude}$, where Maximum E_overshoot_ is the expression level difference between the recovery and baseline timepoints for each gene, and E_amplitude_ is the expression level difference between the postflight and baseline timepoints. In our data, if the gene is significantly upregulated:

$$overshoot=(E_{baseline} - E_{recovery} )/( E_{postflight} - E_{baseline})$$

whereas if the gene is significantly downregulated:

$$overshoot=(E_{recovery}- E_{baseline})/(E_{baseline}-E_{postflight})$$

The mean overshoot value for 90-day flight (M90) was greater than that for180-day flight (M180-1), namely, 0.237 for upregulated genes and 0.209 for downregulated genes; and 0.100 for upregulated genes and 0.144 for downregulated genes, respectively. In practice, such overshoot is observed in oscillating systems. In second-order systems, the percent overshoot is the percentage, by which a system's step response exceeds its final steady-state value, which is related to the damping ratio by the following equation:

$$\zeta=\frac{-ln(percent overshoot)}{\sqrt[2]{\pi^{2}+{ln}^{2}(percent overshoot)}}$$

Here, the percent overshoot can be presented as the mean of the overshoot quantification for both up- and downregulated genes. The damping ratios calculated with the above equation are 0.416 and 0.592 for M90 and M180-1, respectively. To estimate the settling time, we can use the following equation:

$$T_{s}=\frac{-\ln(tolerance fraction)}{\zeta\omega_{n}}$$

In this equation, $\omega_{n}$ is the natural frequency at which the system oscillates when the damping ratio is zero. As the transcriptome of an organism can be considered a dynamic system that is regulated by many molecular mechanisms such as transcriptional control, post-transcriptional control, epigenetic control, and others, the difficulty of determining the natural frequency cannot be ignored. However, the intrinsic number of DEGs can be expressed as:

$$T_{s}=\frac{-\ln(tolerance fraction)}{\zeta\frac{1}{DEG number}}$$

In this way, we are presented with the difficult quantification of the settling time, which we cannot assign a unit. The medium-duration spaceflight has a Ts of 742.093, and the long-duration spaceflight Ts is 818.570. In this scope, the recovery from medium-duration and long-duration spaceflights manifests differently at transcriptomic level. The gene expression recovery from the medium-duration spaceflight appears to be more oscillatory and efficient than of the long-duration spaceflight. This discovery is consistent with reality, and the underlying mechanism is intriguing.

#### Note S3. Bone loss and coagulation activation in HLU model

The bone morphology images intuitively demonstrate a decrease in bone mass in the 28-day HLU model rats. Additional evidence also points to these rats having less bone mass, as seen by a decrease in trabecular number (Tb.N) (FC=0.633, *P*=0.042, Wilcox.test) and trabecular thickness (Tb.Th) (FC=0.828, *P*=0.024, Wilcox.test), alongside an increase in trabecular separation (Tb.Sp) (FC=2.040, *P*=0.042, Wilcox.test) (Table S7). This is typically observed when bone resorption exceeds bone formation, as seen in osteoporosis. The decrease in bone volume fraction (BV/TV) also supports reduced bone mass (FC=0.519, *P*=0.006, Wilcox.test), reflecting the amount of trabecular bone within the medullary cavity. The aforementioned indices are traditionally indices (TRI)^6^ derived from the primary indices assuming a constant structure model and applying stereological techniques. Additionally, the decrease in trabecular connectivity density (FC=0.470, *P*=0.042, Wilcox.test) further indicates bone loss.

Elevated levels of P-selectin (FC=1.505, *P*=0.343, Wilcox.test) and PF4 (FC=1.569, *P*=0.057, Wilcox.test) were detected in the blood of rats exposed to the 28-day HLU model, indicating activation of coagulation.

### Supplemental Tables

#### Table S1.

Detailed information of sampling times for each dataset in this study.

| **Experiments** | **Measurements** | **Subjects** | **Time points** | **Sampling time points** | **Species** |
| --- | --- | --- | --- | --- | --- |
| 13-day spaceflight M13 | DNA methylation;  Phenotypes | 3 | 4 | Pre50 Post1 Post10 Post30 | Human |
| 15-day spaceflight M15 | DNA methylation; Phenotypes | 3 | 4 | Pre50 Post1 Post10 Post30 | Human |
| 33-day spaceflight M33 | DNA methylation; Phenotypes | 2 | 4 | Pre50 Post1 Post10 Post30 | Human |
| 90-day spaceflight M90 | DNA methylation; RNA-seq;  Proteomics;  Protein acetylation;  Phenotypes | 3 | 3 | Pre57 Post1 Post63 | Human |
| 180-day spaceflight M180-1 | DNA methylation; RNA-seq;  Proteomics;  Protein acetylation;  Phenotypes | 3 | 3 | Pre174 Post1 Post68 | Human |
| 180-day spaceflight M180-2 | DNA methylation; RNA-seq | 3 | 4 | Pre48 Post2 Post67 | Human |
| 90-day HDBR | DNA methylation; Phenotypes | 8 | 5 | Pre15 R30 R60 R90 Post30 | Human |
| 180-day CELSS | DNA methylation;  Transcriptome Array;  Phenotypes | 4 | 11 | Pre30 R2 R30 R60 R75 R90 R105 R120 R150 R175 Post30 | Human |
| Mars500 | DNA methylation;  Phenotypes | 6 | 10 | Pre7 R60 R120 R168 R249 R300 R360 R418 R512 Post7 | Human |
| 28-day hindlimb  unloading (HLU) model | RNA-seq;  Phenotypes | 4 (case)  vs. 4 (control) | 2 | D28 (case) vs.  D0 (control) | Rat |
| 35-day hindlimb  unloading (HLU) model | RNA-seq | 3 (case)  vs. 3 (control) | 2 | D35 (case) vs.  D0 (control) | Rat |

#### Table S2.

Results of quantitative and functional enrichment analysis of DMRs

#### Table S3.

Differential expression gene analysis results

#### Table S4.

Differentially spliced genes

#### Table S5.

Differential expression analysis results of peptides and protein acetylation modification changes in mission M90 and M180-1

#### Table S6.

PPI network

#### Table S7.

HLU model rat
